# Unveiling the Hidden Rules: Enhancing NMD Prediction for Protein-Truncating Variants

**DOI:** 10.64898/2026.06.26.734884

**Authors:** Iman Egab, Jacob Schmidt, Michael Cortázar, Jiaoyang Xu, Peter Orchard, Moez Dawood, Tugce Bozkurt-Yozgatli, Jasmine Koh, Luisa Mestroni, Matthew Taylor, S. Stephen Yi, Daniel Calame, Jennifer E. Posey, Richard A. Gibbs, Eric Boerwinkle, Alexander P. Reiner, Paul S. de Vries, Alanna C. Morrison, Chad A. Shaw, James R. Lupski, Claudia M.B. Carvalho, Stephen Montgomery, Sujatha Jagannathan, Zeynep Coban-Akdemir, NHLBI Trans-Omics for Precision Medicine (TOPMed) Consortium

## Abstract

Nonsense-mediated decay (NMD) is a conserved RNA quality-control pathway that degrades transcripts containing premature termination codons. Because roughly a third of pathogenic variants in ClinVar can lead to truncated protein synthesis, predicting whether such transcripts undergo NMD is central to interpreting variant effects, yet the canonical 50–55 nucleotide rule explains only about half of observed outcome variability. Using paired whole-genome and RNA-sequencing from 10,306 individual samples in the Trans-Omics for Precision Medicine (TOPMed) program, we quantified NMD efficiency for 5,749 germline truncating variants via allele-specific expression and trained a gradient-boosting classifier, TrunCat, that distinguished NMD-sensitive from NMD-escape transcripts with ∼78% ROC-AUC (Receiver Operating Characteristic - Area Under the Curve). A reduced model using eight features selected by their mean SHAP (SHapley Additive exPlanations) value, a measure of each feature’s average contribution to predictions, nearly matched this performance. Applied across large variant databases and a rare-disease cohort, the model produced NMD outcome predictions, with variants of uncertain significance showing higher predicted escape than pathogenic ones. This framework confirms the canonical rule, identifies non-canonical determinants, and offers a scalable resource for interpreting protein-truncating variants.

## Introduction

Protein-truncating variants (PTVs) are major contributors to severe genetic disease and account for much of the diagnostic yield of clinical exome and genome sequencing across congenital, neurodevelopmental, and psychiatric disorders ^1–12^. Nonsense-mediated mRNA decay (NMD) is a conserved RNA-surveillance pathway that degrades transcripts harboring premature termination codons (PTCs), limiting accumulation of deleterious truncated proteins^13–19^. In mammals, NMD is coupled to splicing through the exon junction complex (EJC), deposited ∼20–24 nucleotides upstream of every spliced junction^20^. When translation terminates more than ∼50–55 nucleotides upstream of the last EJC, the ribosome and its release factors recruit the core NMD machinery (UPF1, UPF2, UPF3B, SMG1) through the downstream EJC to commit the transcript to decay^16,17,19,21–28^. This EJC-dependent “50–55 nt rule” anchors most genome-wide NMD prediction tools^29–31^ and the PVS1 loss-of-function criterion in current American College of Medical Genetics and Genomics/Association for Molecular Pathology (ACMG/AMP) guidelines^32,33^.

Despite its utility, the canonical rule predicts NMD outcome only incompletely. In the Geuvadis lymphoblastoid panel, Rivas et al. and Lappalainen et al. found that many predicted PTVs escape NMD and that the rule alone explains only about half of the variability in allele-specific expression (ASE) at heterozygous PTC sites^34–37^. Cancer-genome analyses across thousands of tumors added positional rules including reduced efficiency for PTCs in the first ∼150 nt, in long middle exons (>400 bp), and within ∼55 nt of the last EJC and an integrated model explaining roughly three-quarters of non-random variance in NMD efficiency^37–39^; gene-level intolerance to loss-of-function variation is itself a strong predictor^40,41^. Biochemical and single-molecule studies have added non-canonical determinants: long three prime untranslated regions (3ʹUTRs) recruit UPF1 to trigger EJC-independent decay^41–45^; 3ʹUTR composition, GC content, and conserved elements modulate UPF1 deposition and turnover^45–48^; reinitiation downstream of 5ʹ-proximal PTCs can bypass NMD^49–54^; and the rate of peptidyl-tRNA hydrolysis, set by the codon immediately upstream of the PTC, tunes efficiency per substrate^55^.

NMD activity also varies across tissues, individuals, and disease contexts: tissues with defective machinery show globally aberrant splicing^56^, and population-scale, tissue-resolved analyses reveal variation between tissues, individuals, and tumor versus matched-normal samples^57,58^. This has clinical consequences: NMD can aggravate or alleviate disease depending on whether the truncated product is deleterious (NMD-protective) or partially functional (NMD-aggravating), and the same variant may behave differently across backgrounds^45,58–62^. These all make NMD a therapeutic target: its inhibition is being pursued to rescue truncated-but-functional protein and to enhance immune recognition of tumor neoantigens, with small-molecule inhibitors and readthrough agents in clinical development^20,32,59,62–67^

Translating these insights into germline variant interpretation remains challenging. Rule-based annotators including a framework for genes enriched in NMD-escape alleles acting through dominant-negative or gain-of-function (DN/GoF) mechanisms^68^, the aenmd R/Bioconductor package^69^, and the earlier NMD Classifier^30^ mainly encode the 50–55 nt rule with a few hand-curated escape heuristics. Recent machine-learning tools integrate broader signals: Iha et al. combined multi-nucleotide variant rescue, translational status, and isoform expression^29^; NMDEP couples sequence embeddings with curated features^70^; and NMDetective-AI is an end-to-end model built on the Orthrus mRNA foundation model and deep mutational scanning data^31^. Still, few jointly model the transcript-architecture, mRNA stability, conservation, and sequence context features now known to shape NMD, or return per-variant probabilities suitable for clinical use.

These gaps are visible in clinical databases. About a third of pathogenic or likely-pathogenic ClinVar variants are protein-truncating, yet many PTC-variants, even in well-established disease genes, remain variants of uncertain significance (VUS)^32,33^, and PTC-bearing VUS are enriched for predicted NMD-escape relative to high-confidence pathogenic PTCs^33,70^. Regional-constraint analyses of 730,947 individuals show that NMD-informed metrics improve loss-of-function interpretation, with *de novo* PTCs in NMD-constrained regions enriched up to 9.5-fold in rare-disease cohorts^41^. Functional reclassification works but is slow and costly: In one effort ∼75% of 400 VUS were reclassified as pathogenic after targeted assays^6^ underscoring the need for scalable, mechanistically grounded models that resolve ambiguity and prioritize variants for validation.

To address this gap, we present a large-scale empirical and machine-learning analysis of germline PTC-variants. Using paired whole-genome (WGS) and RNA sequencing (RNA-Seq) from 10,306 individual samples in the NHLBI Trans-Omics for Precision Medicine (TOPMed) program, we quantified NMD efficiency by ASE at 5,749 heterozygous nonsense variants in 3,290 canonical transcripts, characterized canonical and non-canonical determinants of NMD across variant-, transcript-, and sequence-context features, and trained an interpretable gradient-boosting classifier (TrunCat, TRUNcation-aware Classifier using Annotated Transcripts; built on CatBoost). We applied TrunCat to four external sets including the Genome Aggregation Database (gnomAD)^71,72^, ClinVar pathogenic/likely-pathogenic (P/LP) variants^73^, ClinVar VUS, and a rare-disease cohort from the Genomic Research Elucidates the Genetics of Rare Disease (GREGoR) consortium to generate predictions, benchmarked against prior cancer-genome and disease-wide analyses of NMD^29,31,37,38,70,73^.

## Materials and Methods

### Data Sources

#### TOPMed

The study population consists of individuals with matched WGS and RNA-Seq data available through the National Heart, Lung, and Blood Institute (NHLBI) Trans-Omics for Precision Medicine (TOPMed) program. The TOPMed Freeze 9b dataset is a refined version of freeze 9. This freeze 9b dataset consists of a total of ∼160.8K samples including: ∼158.3K TOPMed samples and 2,504 samples from 1000 Genome project (https://www.internationalgenome.org/1000-genomes-summary/).

#### The Genome Aggregation Database version 4.1 (gnomAD v4.1) (https://gnomad.broadinstitute.org/)

A large reference resource aggregating human genetic variation across diverse populations, providing allele frequency and constraint metrics from exome and genome sequencing data.

#### ClinVar (release clinvar_20260201) (https://www.ncbi.nlm.nih.gov/clinvar/docs/downloads/)

A publicly available archive of clinically annotated variants and their relationships to human disease phenotypes.

#### Genomic Research Elucidates the Genetics of Rare Disease (GREGoR) (https://gregorconsortium.org/data)

A rare disease cohort resource providing genomic and phenotypic data to support variant pathogenicity assessment and novel disease gene discovery.

### RNA-seq Data Generation and Processing

RNA-seq data for the matched ASE cohort were generated within the contributing TOPMed studies including the Framingham Heart Study (FHS), Gene-Environments and Admixture in Latino Asthmatics (GALA II), Study of African Americans, Asthma, Genes & Environments (SAGE), Subpopulations and Intermediate Outcome Measures in COPD Study (SPIROMICS), Women’s Health Initiative (WHI), COPDGene, Multi-Ethnic Study of Atherosclerosis (MESA), and Lung Tissue Research Consortium (LTRC) across six tissues and cell types: whole blood, lung, peripheral blood mononuclear cells (PBMC), monocytes, T cells, and nasal epithelium^74,75^. The samples retained for ASE analysis were drawn predominantly from whole blood (n = 6,602), with additional contributions from lung (n = 1,360), PBMC (n = 1,265), T cell (n = 368), nasal epithelial (n = 359), and monocyte (n = 352) samples (10,306 RNA-seq samples in total) (**Table S1**); whole blood was contributed by multiple cohorts, whereas each remaining tissue derived from a single cohort^75^.

For blood-derived samples, total RNA was extracted from PAXgene-stabilized whole blood or from isolated cell populations, while lung and nasal-epithelial RNA was extracted from the corresponding tissues. Strand-specific libraries were prepared following poly-A selection using the Illumina TruSeq Stranded mRNA kit and sequenced at the Northwest Genomics Center and the Broad Institute on Illumina HiSeq 4000 or NovaSeq 6000 instruments^75^.

Reads were aligned to the GRCh38 reference genome including ERCC spike-ins and excluding ALT, HLA, and decoy contigs using STAR^76^ with a collapsed GENCODE v30 gene model produced by the GTEx collapse-annotation pipeline^77,78^; MESA samples were aligned with an earlier STAR build indexed on GENCODE v26. Duplicate reads were marked with Picard, and gene-level read counts, transcript-per-million (TPM) values, and per-sample quality-control metrics were computed with RNA-SeQC 2^79^. Outlier samples were identified from a principal-component analysis of normalized expression and excluded from downstream analysis^75^.

Each RNA-seq sample was matched to its TOPMed Freeze 9b whole-genome genotypes^74^ by calling approximate genotypes directly from the RNA alignments (restricted to reads with mapping quality ≥ 20) and computing non-reference genotype concordance against all Freeze 9b samples at PASS coding-exon variants with TOPMed-wide minor allele frequency ≥ 0.05; a sample was considered matched when its highest genotype concordance exceeded 83%^74,75^. To eliminate reference-allele mapping bias prior to allele-specific counting, reads were re-aligned with STAR in a variant-aware manner using WASP filtering, and only uniquely mapped reads passing the WASP filter were retained^80,81^. These bias-corrected, genotype-matched alignments provided the input for the allele-specific read counting described below.

### Dataset Sources

#### TOPMed

Biallelic variants were extracted from TOPMed Freeze 9b whole-genome sequencing (WGS) files (BCF/VCF format) spanning chromosomes 1–22 and X. 10,306 individual samples with both high-quality matched RNA-Seq and WGS data was selected for integrated ASE analysis. Variants were annotated using SnpEff^83^ (https://pcingola.github.io/SnpEff/snpeff/inputoutput/), and those predicted to introduce premature termination codons (PTCs) including nonsense and frameshift variants were selected using *bcftools.* This yielded 77,918 candidate variants. To ensure compatibility with downstream ASE quantification, variants with overlapping genomic positions were removed, resulting in a final set of 77,408 variants. ASE was then quantified at these sites using GATK ASEReadCounter (v3)^80,82^ (https://gatk.broadinstitute.org/hc/en-us/articles/360037428291-ASEReadCounter), which computes allele-specific read counts from RNA-seq alignments.

Additionally, genotype information was then merged with ASE read count data based on genomic coordinates to generate variant-level tables containing reference and alternate allele counts alongside corresponding genotype calls. Only variants passing standard quality filters (FILTER = PASS) were retained.

#### GnomAD, ClinVar, and GREGoR

Variants from gnomAD, ClinVar and GREGoR were preprocessed to remove duplicate and low-quality entries. Unique variants were annotated for their PTC-generations outcomes using the aenmd (R-package)^69^. Only variants that are PTC-generating were selected for the analysis.

### Allele-Specific Expression and NMD Efficiency Quantification

ASE were quantified for putative PTVs in the TOPMed Freeze 9b dataset comprising 10,306 individual samples with matched RNA-Seq data. Variants were annotated using SnpEff^83^ (https://pcingola.github.io/SnpEff/snpeff/inputoutput/), and those predicted to introduce premature protein truncation were selected using *bcftools.* This yielded 77,918 candidate variants. To ensure compatibility with downstream ASE quantification, variants with overlapping genomic positions were removed, resulting in a final set of 77,408 variants. ASE was then quantified at these sites using GATK ASEReadCounter (v3)^80,82^ (https://gatk.broadinstitute.org/hc/en-us/articles/360037428291-ASEReadCounter), which computes allele-specific read counts from RNA-seq alignments.

To obtain a single allele-specific expression estimate per variant, we implemented a resampling-based approach to account for variants observed across multiple individuals. Variants were first filtered to retain sites with sufficient read depth (total read count ≥ 8). Variants observed in only one individual were treated as singletons, and ASE was calculated directly from the observed reference and total read counts^34^. For variants observed in multiple individuals, a resampling procedure was applied to avoid bias introduced by arbitrary selection of a single observation. Specifically, for each variant, one observation was randomly sampled from the set of available individuals, and the corresponding allelic ratio (reference allele count divided by total read count) was computed. This process was repeated 100 times, generating a distribution of allele ratios per variant. The median allelic ratio across iterations was then used as the final ASE estimate for that variant.

For NMD-sensitive variants, the expected proportion of wild-type allele expression is larger than 0.65; for NMD-escaping variants, the expected allele ratio is between 0.35 and 0.65^84^.

### Gene and Transcript Filtering

To minimize confounding effects and ensure robust estimation of NMD efficiency, stringent filtering criteria were applied to the set of unique variants. First, variants located in single-exon genes were excluded (exon_count > 1). Second, lowly expressed genes in whole blood were removed using median gene-level expression estimates from GTEx Analysis v8 transcripts per million (TPM)^77^ (https://www.gtexportal.org/home/downloads/adult-gtex/bulk_tissue_expression). Third, genes exhibiting significant expression quantitative trait loci (eQTL) effects were excluded based on GTEx v8 eGenes (whole blood; q ≤ 0.05) (https://gtexportal.org/home/downloads/adult-gtex/qtl). Finally, imprinted genes were removed using curated imprinted gene lists provided in **Table S2**.

### Variant Annotation and Transcript Feature Extraction

All PTC-generating variants from TOPMed, GnomAD, ClinVar and GREGoR were first annotated using ANNOVAR^85^ with the Ensembl gene model (ensGene) for GRCh38, based on a database release available in July 2017. ANNOVAR was used to assign functional consequences and extract transcript-level annotations, including gene identity, exon structure, and coding sequence context.

Because ANNOVAR may return annotations for multiple transcripts per variant, canonical transcript IDs were obtained separately from Ensembl BioMart^86^ (https://www.ensembl.org/biomart). For each variant, the ANNOVAR transcript annotation field was parsed, and the annotation corresponding to the BioMart-defined canonical transcript for that gene was retained for downstream feature extraction.

Variant-level functional scores and population frequency metrics were then incorporated using ANNOVAR, including REVEL^87^, CADD Phred-scaled scores^88^, and allele frequencies from gnomAD exomes^71^.

### Genomic Feature Retrieval

Transcript structure annotations were derived from GENCODE v26^78^ (https://www.gencodegenes.org/human/release_26.html), and reference genome sequences were obtained from BSgenome.Hsapiens.UCSC.hg38. These annotations captured key aspects of gene architecture, including coding sequence (CDS) length, exon structure, and exon count, as well as untranslated region (UTR) properties such as 5′ and 3′ UTR length and the presence of upstream open reading frames.

Sequence composition was further quantified across transcripts by calculating dinucleotide content (e.g., AU, GC, UC) across full regions and within defined windows, including the first and last 100–200 base pairs of coding sequences and UTRs. In addition, positional features were derived to describe the location of each PTC within the transcript, including distances to the start and stop codons, exon boundaries, and downstream in-frame AUG codons.

To further capture local sequence context, amino acid features were annotated at positions −2, −1, +1, and +2 relative to the PTC. Coding sequences were translated into protein sequences, and the position of each PTC was mapped onto the corresponding protein to extract flanking residues. These features were motivated by prior work demonstrating that local sequence context influences translation termination efficiency and NMD^57^.

### Gene Constraint Metrics

Gene-level intolerance to loss-of-function variation was assessed using gnomAD v2.1.1 constraint metrics^71,72^ (https://gnomad.broadinstitute.org/downloads). The probability of loss-of-function intolerance (pLI) was used to classify genes as highly tolerant (pLI < 0.35), moderately tolerant (0.35 ≤ pLI < 0.65), or highly intolerant (pLI ≥ 0.65). The loss-of-function observed/expected upper bound fraction (LOEUF) was also included as a continuous measure of constraint, representing a conservative estimate of loss-of-function depletion based on the upper bound of a Poisson-derived confidence interval. Lower LOEUF values indicate stronger purifying selection against predicted loss-of-function variants.

### RNA-Binding Protein Motif Identification

To capture regulatory elements influencing NMD, multiple biologically relevant sequence windows were defined for motif scanning, including: ±100 base pairs surrounding the PTC; the region between the PTC and the nearest downstream EJC; ±100 base pairs surrounding EJCs; the first 200 base pairs and the full length of the 3′UTR; the first 200bp and the full length of the new 3’UTR which defined as the length from the PTC to the end of 3′UTR. Sequences were scanned for known RNA-binding protein (RBP) motifs using FIMO^89^ (MEME Suite version 5.5.5: https://meme-suite.org/meme/doc/download.html), with motif definitions obtained from the ATtRACT database (Homo sapiens:https://attract.cnic.es/)^90^. Binary indicators of motif presence or absence were generated for each region and motif combination.

### Transcript-Level Features

Transcript stability was quantified using PC1-derived mRNA half-life estimates from Agarwal and Kelley^89^, representing a consensus measure across 39 human datasets. This metric is highly correlated (r > 0.99) with mean measured half-life and provides a cell-type-independent proxy for intrinsic mRNA stability. Additionally, gene expression features were incorporated using GTEx version 8^77^ (https://www.gtexportal.org/home/downloads/adult-gtex/bulk_tissue_expression). Median expression across tissues was calculated for each gene, and whole blood expression was retained as a tissue-relevant measure. Expression values were log-transformed prior to analysis.

### EJC Occupancy Quantification

To assess EJC occupancy near premature termination codons, we implemented a transcript-aware windowing approach using experimentally validated EJC binding sites from RNA Immunoprecipitation Sequencing (RIP-Seq) data (https://www.ncbi.nlm.nih.gov/geo/query/acc.cgi?acc=GSE41154). Each variant was analyzed using its canonical transcript as specified earlier in the Methods section. For each PTC, we identified the nearest downstream exon-exon junction within the same exon by querying Gencode v26 transcript annotations and determining strand-specific junction positions. We then computed a 30-nucleotide window centered at −24 nucleotides upstream of this junction (±15 bp), representing the canonical EJC deposition site. EJC occupancy was quantified by overlapping this window with experimentally determined EJC peaks and counting the number of overlapping intervals. PTCs occurring in terminal exons typically yielded zero EJC occupancy due to the absence of downstream junctions, though a small number of terminal exon variants showed non-zero counts, likely reflecting residual EJC signal or annotation edge cases. The analysis was parallelized across chromosomes using Python’s ProcessPoolExecutor, processing variants in batches to optimize memory usage. Both continuous count values and binary indicators (presence/absence of EJC overlap) were retained as features for model training.

### Codon Optimality Scoring

We quantified codon optimality as a measure of translational efficiency following the methodology of Lindeboom et al. (2016)^37^. Optimal codons were defined as those corresponding to tRNA genes with the highest copy number in the human genome, derived from UCSC tRNA gene annotations. For each tRNA gene, we computed the corresponding codon by reverse complementing the anticodon sequence. We then counted tRNA gene copies per codon and selected the codon(s) with maximum representation for each amino acid as “optimal.”

For each variant, we extracted the complete coding sequence (CDS) from Gencode v26 annotations using the canonical transcript specified in the txnames column. CDS sequences were constructed by concatenating CDS segments in coding order and applying reverse complementation for minus-strand transcripts. We computed two codon optimality metrics: (1) whole-CDS optimality fraction, calculated as the proportion of optimal codons across the entire coding sequence, and (2) local optimality fraction within a ±100-nucleotide window surrounding the PTC position. To map genomic PTC coordinates to CDS positions, we converted 1-based VCF-style positions to 0-based coding frame offsets accounting for splicing patterns and strand orientation. Reference genome sequences were extracted from hg38 using pyfaidx, and only complete codons (3-nucleotide units without ambiguous bases) were included in calculations. The analysis was parallelized across variant batches (chunk size: 50,000) using multi-process execution, with CDS sequences cached to avoid redundant genome queries.

### Stop Codon Readthrough Potential

We implemented a context-dependent scoring system to predict translational readthrough probability based on the HEK293T experimental framework from Mangkalaphiban et al. (2024)^91^. The scoring algorithm evaluates the 12-nucleotide sequence context surrounding each stop codon (3 nucleotides upstream through 9 nucleotides downstream) and assigns weighted scores based on experimentally validated features. Stop codon identity contributes 10–40 points (TGA: 40, TAG: 25, TAA: 10), the +4 position contributes up to 20 points (C > T > A/G), and positions +5 through +9 contribute 2–6 points each with stop-codon-specific preferences. The P-site position (−3) contributes 6 points if thymine, otherwise 2 points. Raw scores are normalized to a 0–100 scale and categorized as high (≥70), medium (50–69), low (30–49), or none (<30) readthrough potential.

### Performance Evaluation and Interpretability

We trained the NMD-outcome classifier with CatBoost, a gradient-boosting machine-learning algorithm that combines a series of decision trees, with each successive tree learning to improve the prediction errors of the preceding trees. Model performance was assessed using 5-fold stratified cross-validation (stratified by NMD-outcome class; random_state = 42). Hyperparameters were tuned via Optuna using a TPE sampler with 100 trials, where each trial’s objective was mean ROC-AUC across the same 5-fold CV split; the search covered iterations, learning_rate, depth, l2_leaf_reg, bagging_temperature, random_strength, border_count, and min_data_in_leaf, with early stopping (50 rounds) on each fold’s validation set. We acknowledge that hyperparameter tuning and performance evaluation share the same 5-fold split, which can introduce mild optimism in reported AUC. The final model was refit using the best hyperparameters and evaluated by concatenating predictions across the five validation folds to obtain a single set of out-of-fold (OOF) predictions covering all training variants. Reported ROC-AUC and PR-AUC are computed on these OOF predictions, with 95% confidence intervals obtained by bootstrap resampling (1,000 iterations). The Youden-optimal classification threshold (τ = 0.47) was selected on the concatenated OOF predictions; we recognize that selecting the threshold on the same predictions used to evaluate threshold-dependent metrics (sensitivity, specificity, accuracy) introduces a small additional optimism, and we accordingly emphasize threshold-free metrics (ROC-AUC, PR-AUC) in our headline reporting. Feature importance was quantified using SHAP (TreeExplainer) on the final CatBoost model. Feature-ablation analysis re-fit CatBoost models with the top-k features ranked by mean |SHAP| value at each step, with each top-k model independently evaluated under the same 5-fold stratified CV.

### External Application

The trained final model was used to generate per-variant NMD-escape predictions on to four external datasets: (i) all nonsense variants in gnomAD v2.1.1^71^ (n = 137,857); (ii) ClinVar pathogenic/likely-pathogenic nonsense variants^73^(P/LP; n = 23,856); (iii) ClinVar variants of uncertain significance (VUS; n = 3,445); and (iv) nonsense variants identified in probands enrolled in the GREGoR (Genomics Research to Elucidate the Genetics of Rare Diseases) consortium (n=2,548). Features were extracted using the same pipeline as TOPMed variants. Predicted-escape probabilities were thresholded at the Youden-optimal cutoff (τ = 0.47) to assign NMD-sensitive vs. NMD-escape calls.

### Experimental Validation Using LMNA Saturation Genome Editing

To evaluate concordance between model predictions and experimentally measured NMD activity, we applied TrunKitten to the LMNA saturation genome editing (SGE) dataset reported by Cortázar et al. (2025)^92^, who systematically introduced PTCs across the first, internal, and penultimate exons of LMNA and quantified NMD efficiency by comparing variant transcript abundance with and without pharmacological NMD inhibition. Because TrunKitten was developed for single-nucleotide stop-gain variants, we restricted the analysis to engineered PTCs that could be generated from the endogenous LMNA codon by a single-nucleotide substitution; of 770 PTC measurements, 86 met this criterion. The eight TrunKitten features were reconstructed for each variant from the canonical LMNA transcript (ENST00000368300) using the same feature definitions as in model development, and the pretrained TrunKitten model was applied without retraining. Predicted NMD probability was calculated as 1 − escape probability, and variants were classified as NMD-escape or NMD-triggering at TrunKitten’s decision threshold (escape probability 0.43). Experimental NMD efficiencies were normalized by subtracting the median NMD efficiency of SNV controls within the corresponding SGE region, and variants with an experimental standard deviation > 0.2 were excluded, leaving 84 variants. Concordance was assessed with Spearman rank correlation, and experimental NMD efficiency was compared between predicted classes with a two-sided Wilcoxon rank-sum test.

### Software and Computational Environment

Analyses were performed using Python 3.12.7 with CatBoost 1.2.8, scikit-learn 1.5.1, SHAP 0.48.0, Optuna 4.5.0, pandas 2.2.2, and numpy 1.26.4. All the code and annotations are available at (https://github.com/CobanAkdemirlab/NMDpredictionmodel).

## Results

### Population-Scale Allele-Specific Expression Quantifies NMD Efficiency for 5,749 Nonsense Variants

We analyzed paired RNA-seq and WGS data from the NHLBI TOPMed cohort to identify features associated with NMD outcomes (**Figure 1**). ASE was quantified across 10,306 individual samples to infer NMD efficiency for heterozygous PTC-variants, retaining 5,749 high-quality nonsense variants in 3,214 different canonical transcripts after filtering. To minimize confounding effects and ensure robust NMD efficiency estimation, stringent filtering criteria were applied. Specifically, we excluded: (i) variants in single-exon genes (exon_count > 1), (ii) lowly expressed genes in whole blood based on GTEx v8 median transcript per million (TPM) values, (iii) genes with significant expression quantitative trait loci (eQTL) effects using GTEx v8 eGenes (whole blood; q ≤ 0.05), and (iv) imprinted genes based on curated gene lists

**Figure 1.**
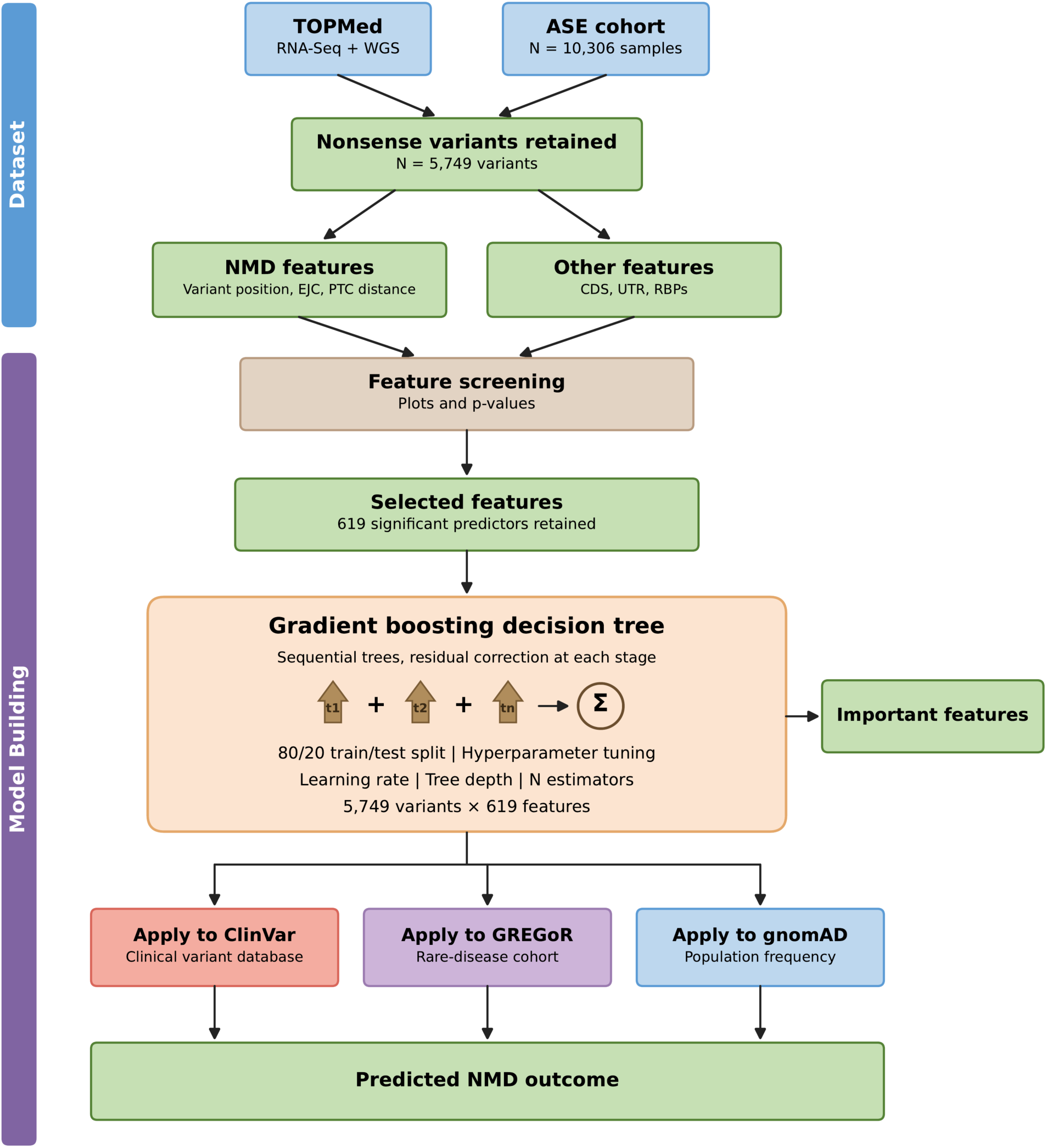
Overview of the analytical and model-building workflow. Paired RNA sequencing (RNA-Seq) and whole genome sequencing data from the Trans-Omics for Precision Medicine (TOPMed) cohort (n = 10,306 individual samples available with allele-specific expression (ASE cohort)) were used to identify 5,749 high-quality nonsense variants. Variant- and transcript-level features (NMD-related: variant position, exon junction complex (EJC), premature termination codon (PTC) distance; other: coding sequence (CDS), untranslated region (UTR), RNA binding protein (RBP) motifs) were screened by univariate analysis, retaining 619 significant predictors. A CatBoost classifier was trained on the 5,749 × 619 feature matrix using five-fold gene-grouped cross-validation with Optuna hyperparameter tuning. The trained model was applied to ClinVar, the Genomic Research Elucidates the Genetics of Rare Disease (GREGoR), and the Genome Aggregation Database (gnomAD) to generate NMD-escape vs. NMD-sensitive predictions and feature importance rankings.

Candidate NMD features (EJC position, variant position, PTC distance) and other transcript-level features (CDS, UTR, and RBP motifs) were evaluated by univariate plots and statistical tests, yielding 619 retained predictors. A machine-learning model (CatBoost), which combines a series of decision trees to make predictions, was trained on the 5,749 variants and their 619 features to distinguish NMD-sensitive from NMD-escape variants. Model performance was evaluated using five-fold cross-validation: the data were divided into five groups, with all variants from the same gene kept in the same group, and the model was trained on four groups and tested on the remaining one, so that each group was used for testing once. Model settings were optimized using Optuna (see Methods). The trained model was then applied to four external variant sets: ClinVar (clinical variant database including P/LP and VUS separately), GREGoR (rare-disease cohort), and gnomAD (population frequency), to generate predicted NMD-escape vs. NMD-sensitive calls and ranked feature importance for downstream clinical and mechanistic interpretation.

### An Interpretable Gradient-Boosting Model, TrunCat, Predicts NMD Outcomes and Identifies a Compact Feature Set

We trained TrunCat, a gradient-boosting classifier built on CatBoost, on the 5,749 × 619 feature matrix to predict NMD outcome from variant- and transcript-level context using five-fold gene-grouped cross-validation (**Figure 2**; see Methods).

**Figure 2.**
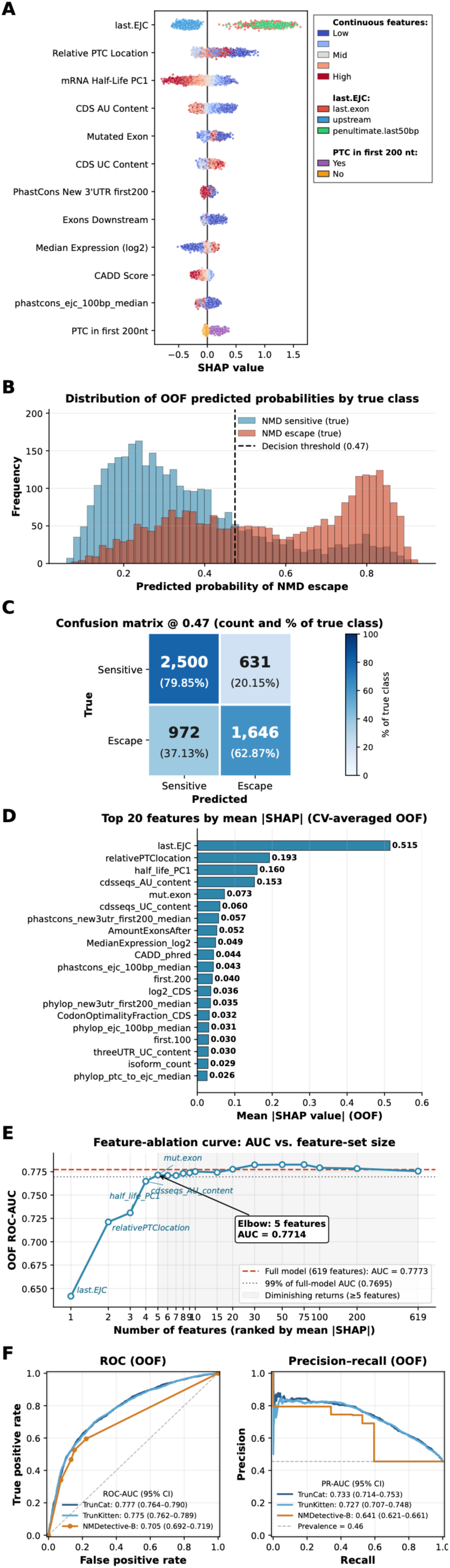
Performance and interpretation of the CatBoost-based TrunCat NMD-outcome prediction model. (A) SHAP value distributions for the 12 most influential features, colored by feature value (blue–red gradient for continuous features; categorical for last.EJC and PTC-in-first-200-nt). (B) Distribution of out-of-fold (OOF) predicted probabilities by true NMD-sensitive vs. NMD-escape class; dashed line marks the decision threshold (τ = 0.47). (C) Confusion matrix at τ = 0.47, showing counts and percentages of the true class: 79.9% of true NMD-sensitive and 62.9% of true NMD-escape variants are correctly classified. (D) Top-20 feature importance ranking by mean |SHAP| value (cross-validation-averaged out-of-fold SHAP values). (E) Feature-ablation curve showing OOF ROC-AUC as features are added in decreasing |SHAP| order; the five-feature elbow (AUC = 0.7714) retains 99% of the full 619-feature model AUC of 0.7773; the gray region marks the diminishing-returns plateau. (F) Receiver-operating-characteristic (ROC) and precision–recall (PR) curves for NMD-escape prediction in the TOPMed cohort. TrunCat and TrunKitten curves are derived from out-of-fold predictions, whereas NMDetective-B is a deterministic, rule-based classifier and is therefore shown as a step function reflecting its discrete output categories. ROC-AUC: TrunCat 0.777 (95% CI 0.764–0.790), TrunKitten 0.775 (0.762–0.789), and NMDetective-B 0.705 (0.692–0.719). Corresponding PR-AUC values are 0.733 (0.714–0.753; TrunCat), 0.727 (0.707–0.748; TrunKitten), and 0.641 (0.621–0.661; NMDetective-B). The horizontal dashed line in the PR panel indicates the escape-class prevalence (0.46).

To interpret the model’s behavior we computed SHAP values (TreeExplainer) for each cross-validation model and averaged them to obtain out-of-fold (OOF) SHAP values for every variant. SHAP value distributions across the 12 most influential features (**Figure 2A**) revealed coherent and directionally consistent contributions. The strongest effect came from last-exon and penultimate-last-50 bp status (last.EJC), which shifted predictions toward NMD escape, while the presence of a downstream EJC shifted predictions toward NMD sensitivity. Relative PTC location, mRNA half-life (PC1)^93^, CDS AU content, mutated-exon number, CDS UC content, conservation of the new 3ʹUTR (PhastCons, first 200 nt) and of the region around the downstream EJC (PhastCons, EJC ± 100 bp), number of downstream exons, median expression, and CADD score each contributed systematically. PTC-in-first-200-nt status, a binary indicator, shifted predictions toward NMD escape. The top-20 ranking by mean |SHAP| value (**Figure 2D**) was consistent with this pattern: ‘EJC downstream of PTC’ was by far the most influential predictor (mean |SHAP| = 0.515), followed by relative PTC location (0.193) and mRNA half-life PC1 (0.160). Other highly ranked predictors captured sequence composition, exon and transcript architecture, evolutionary conservation, and gene expression. Canonical positional features, mRNA-stability features, sequence-composition features, and evolutionary conservation metrics were each represented in the top 20, indicating that the model integrates information across distinct biological axes rather than relying on any single class of feature.

Feature-ablation analysis (**Figure 2E**) showed that this multi-axis integration is achieved with a small number of features. Starting from the single most informative feature (last.EJC) and iteratively adding features in decreasing mean |SHAP| order, Out-of-fold ROC-AUC rose steeply from approximately 0.64 (one feature) to 0.72 (adding relativePTClocation), 0.73 (adding half_life_PC1), 0.76 (adding cdsseqs_AU_content), and 0.771 (adding mut.exon). An elbow was reached at five features (Out-of-fold ROC-AUC = 0.771), 99.2% of the full 619-feature model’s OOF ROC-AUC of 0.777; beyond this point, performance changed little (0.775 with 10 features and at most 0.783). Because performance was maintained with well under ten features, we defined a parsimonious model, TrunKitten, using a pre-specified stability rule: a feature was retained only if it ranked among the ten most important features by mean |SHAP| and also ranked in the top ten in each of five leave-one-fold-out rankings. Eight features met this rule including last.EJC, relativePTClocation, half_life_PC1, cdsseqs_AU_content and mut.exon) and we report TrunKitten alongside TrunCat throughout. Additional analyses of feature redundancy, compensation after removal of last.EJC, and alternative TrunKitten feature sets are provided in **Figure S1**.

Out-of-fold predicted-probability distributions showed strong but imperfect separation by class (**Figure 2B**): NMD-sensitive variants concentrated at predicted probabilities <0.45, while NMD-escape variants concentrated at probabilities >0.7. We selected a reference classification threshold of τ = 0.47 by maximizing Youden’s J on these OOF predictions; we note that this threshold was selected on the same predictions used to evaluate threshold-dependent metrics below and so should be regarded as a reference operating point. At τ = 0.47, TrunCat assigned the correct class to 79.9% of NMD-sensitive variants (2,500 of 3,131) and 62.9% of NMD-escape variants (1,646 of 2,618), with confusion-matrix counts of 2,500 true-sensitive, 631 false-escape, 972 false-sensitive, and 1,646 true-escape (**Figure 2C**).

We benchmarked TrunCat against TrunKitten and the rule-based NMDetective-B classifier on the TOPMed cohort using out-of-fold predictions for the two CatBoost models and per-variant rule-based scores for NMDetective-B, with identical canonical-transcript mapping and the same ASE-derived NMD-outcome labels in all three cases **(Figure 2F**; see Methods). TrunCat achieved the highest discrimination by both ROC-AUC (0.777, 95% CI 0.764–0.790) and PR-AUC (0.733, 95% CI 0.714–0.753), essentially indistinguishable from TrunKitten (ROC-AUC 0.775, 95% CI 0.762–0.789; PR-AUC 0.727, 95% CI 0.707–0.748) and above NMDetective-B (ROC-AUC 0.705, 95% CI 0.692–0.719; PR-AUC 0.641, 95% CI 0.621–0.661). The overlapping 95% confidence intervals of TrunCat and TrunKitten reinforce the feature-ablation finding that a small number of well-chosen features captures nearly all of the predictive signal. TrunCat also showed higher discrimination than NMDetective-B (Δ ROC-AUC = 0.072; Δ PR-AUC = 0.092), suggesting that the additional non-canonical, mRNA-stability, and sequence-context features may provide predictive information beyond canonical rule-based features. However, NMDetective-B was originally developed using NMD outcomes defined in a different dataset and experimental context than the ASE-derived outcomes used here. Its performance in the TOPMed cohort therefore reflects how well the published model generalizes to our NMD outcome definition, rather than a direct comparison between models trained and evaluated under identical conditions.

Stratifying the out-of-fold escape probabilities by NMDetective-B rule bin (**Figure S2**) showed that, for both TrunCat and TrunKitten, the median predicted escape probability closely tracked the empirical escape rate within each bin, from highest in the last_exon bin (n = 1,123) to lowest in the trigger_NMD bin (n = 3,493), while the wide spread of predictions within each bin reflected variant-level differences that the rule assignment alone does not capture.

### The Canonical EJC-Based Rule Is Recapitulated and Refined

We next dissected the top-ranked signals identified by the model, beginning with the canonical EJC-dependent rule^16,17,19,22–26^. Across all heterozygous nonsense variants, we observed a clear positional effect consistent with this rule (**Figure 3A**): PTCs in upstream (non-last, non-penultimate exon last 50 bp) exons (n = 4,394) exhibited significantly higher NMD efficiency (median ∼0.73) than those in the last exon (n = 1,123; median ∼0.54) or the penultimate exon last 50 bp (n = 232; median ∼0.56), with both upstream-versus-last-exon and upstream-versus-penultimate comparisons highly significant (Wilcoxon rank-sum test, P < 0.0001), while last-exon and penultimate-exon last 50 bp variants did not differ significantly from each other (ns).

**Figure 3.**
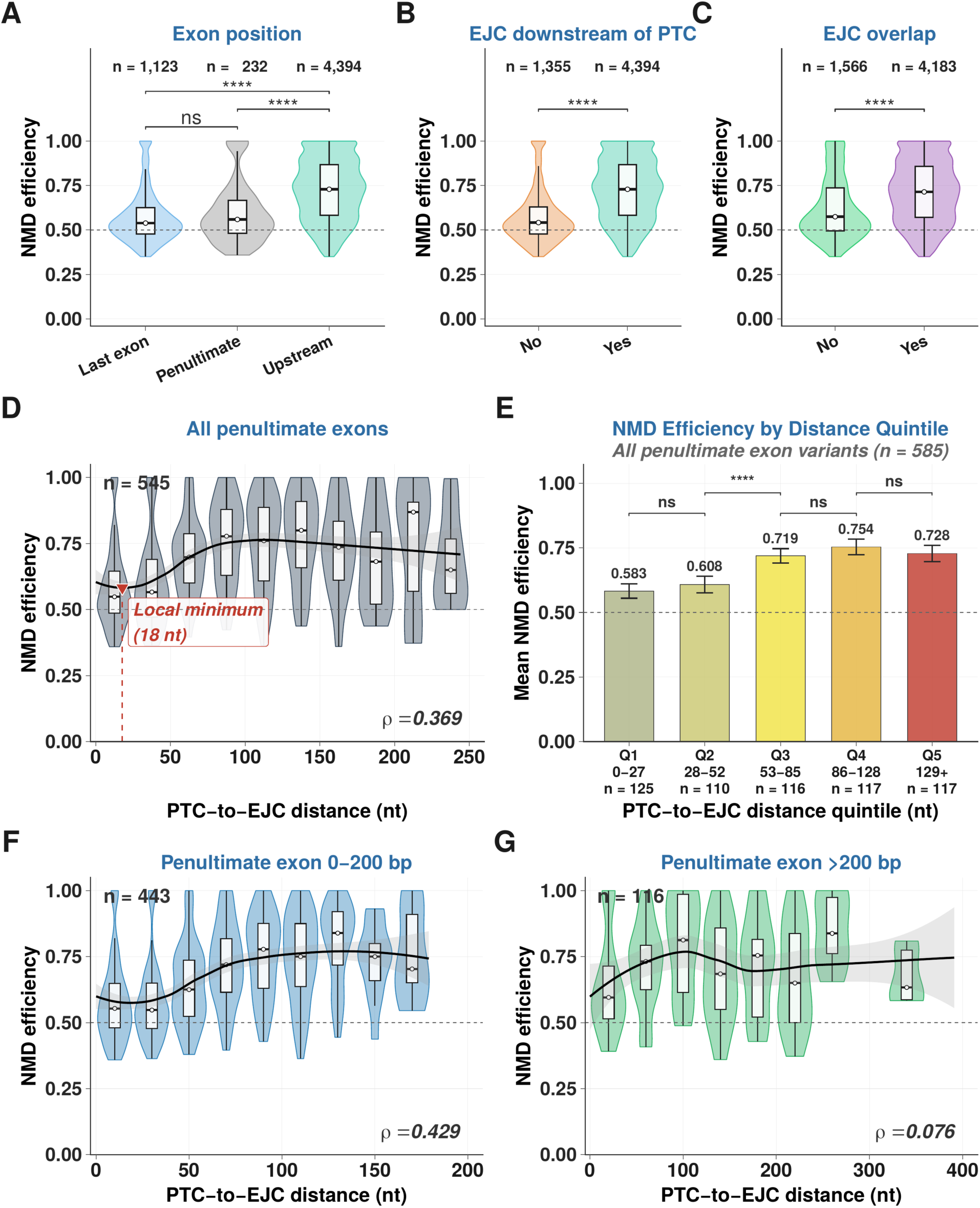
**(A)** NMD efficiency by PTC exon position (last exon n = 1,123, penultimate n = 232, upstream n = 4,394); upstream variants show significantly higher NMD efficiency than both last-exon and penultimate-exon last 50 bp variants (both **** P < 0.0001), while last-exon and penultimate-exon last 50-bp variants do not differ significantly (ns). **(B)** NMD efficiency stratified by presence/absence of a downstream EJC (No n = 1,355, Yes n = 4,394; **** P < 0.0001). **(C)** NMD efficiency stratified by EJC occupancy, defined as overlap between the premature termination codon and experimentally validated EJC binding sites within a 30-nucleotide window centered at −24 nucleotides upstream of the nearest downstream exon–exon junction (No overlap n = 1,566, Overlap detected n = 4,183; **** P < 0.0001). **(D)** PTC-to-EJC distance versus NMD efficiency for penultimate-exon variants in transcripts with ≥ 3 exons and PTC-to-EJC distance ≤ 250 nt (n = 545), shown as violin and box plots per 25-nt distance bin; LOESS smoother (black) with the local minimum annotated at 18 nt; Spearman ρ = 0.369. **(E)** Mean NMD efficiency (± 95% CI) across PTC-to-EJC distance quintiles for all penultimate-exon variants (n = 585; Q1 0–27 nt, Q2 28–52 nt, Q3 53–85 nt, Q4 86–128 nt, Q5 129–4,291 nt; n = 125/110/116/117/117); among adjacent-quintile comparisons, only Q2-vs-Q3 reaches significance (****), while Q1-vs-Q2, Q3-vs-Q4, and Q4-vs-Q5 are ns). **(F, G)** Distance dependence stratified by penultimate-exon length: **(F)** 0–200 bp (PTC-to-EJC distance ≤ 200 nt; n = 443; ρ = 0.429) and **(G)** >200 bp (PTC-to-EJC distance ≤ 400 nt; n = 116; ρ = 0.076). All group comparisons by Wilcoxon rank-sum test (P < 0.05, ** P < 0.01, *** P < 0.001, **** P < 0.0001); correlations are Spearman’s ρ.

Comparing variants with or without a downstream EJC directly tested the canonical 50–55 nt rule (**Figure 3B**): variants with a downstream EJC (n = 4,394) showed markedly higher NMD efficiency (median ∼0.73) than those without (n = 1,355; median ∼0.54; P < 0.0001). EJC occupancy, which flags variants whose PTC falls within the canonical EJC deposition footprint based on the analysis of RIP-Seq data, yielded the same pattern (**Figure 3C**; n = 4,183 with overlap vs. 1,566 without; P < 0.0001) (**Figure 3C**).

To examine the quantitative form of the PTC-to-EJC distance dependence, we restricted our analysis to penultimate-exon variants in transcripts with at least three exons and a PTC-to-EJC distance ≤ 250 nt (n = 545). A non-monotonic relationship emerged (**Figure 3D**; Spearman ρ = 0.369): NMD efficiency was lowest at very short PTC-to-EJC distances, with a local minimum of the fitted LOESS curve near 18 nt, increased across the 20–100 nt window, peaked around 100– 150 nt, and declined modestly at larger distances. In quintile form, using all 585 penultimate-exon variants without a distance cap (**Figure 3E**), mean NMD efficiency changed from 0.583 in Q1 (0– 27 nt; n = 125) and 0.608 in Q2 (28–52 nt; n = 110) to 0.719 in Q3 (53–85 nt; n = 116), 0.754 in Q4 (86–128 nt; n = 117), and 0.728 in Q5 (129–4,291 nt; n = 117); among adjacent quintiles, only the Q2-to-Q3 increase was significant (P < 0.0001).

Stratification by exon length (**Figure 3F, G**) showed that exon architecture strongly modifies the distance dependence. In short-to-medium penultimate exons (0–200 bp; n = 443), the PTC-to-EJC distance effect was strong and well-resolved (Spearman ρ= 0.429), with the LOESS smoother showing a clear monotonic rise from ∼0.55–0.60 at very short distances up to ∼0.75 by 100–150 nt. In long penultimate exons (>200 bp; n = 116), the distance effect was weak (ρ = 0.076) and the LOESS smoother showed only a flat trend across the broader 0–400 nt range, indicating that within long penultimate exons the PTC-to-EJC distance becomes a much weaker determinant of NMD efficiency, consistent with prior reports that long exons partially evade canonical NMD.

### The Canonical Rule Holds Across Gene Constraint, Allele Frequency, and Transcript Architecture

As a baseline, we first examined the marginal distributions of NMD efficiency across these same architectural strata (**Figure S3**). Marginally, gene tolerance (highly intolerant, n = 349; highly tolerant, n = 2,772 transcripts; Mann-Whitney ns) and CDS length (short <1.3 kb, n = 1,302; medium 1.3–2.2 kb, n = 1,029; long >2.2 kb, n = 883; Kruskal-Wallis ns) showed no significant effect on NMD efficiency, allele frequency (ultra-rare, n = 4,165; rare/common, n = 1,584) showed only a small significant difference (P < 0.05), and exon count (2–5, n = 696; 6–10, n = 1,024; 11– 20, n = 1,021; >20, n = 473) showed a strong overall effect (Kruskal-Wallis P < 0.0001) driven primarily by lower NMD efficiency in 2–5-exon transcripts. To confirm that the canonical EJC effect was not confounded by selection, allele-frequency class, or transcript architecture, we then stratified variants by probability of loss-of-function intolerance (pLI) bin, allele frequency class, CDS length tertile, and exon count bin, and compared NMD efficiency between variants with and without a downstream EJC within each stratum (**Figure S4**). The downstream-EJC effect was highly significant in every stratum. In highly loss-of-function intolerant genes (pLI ≥ 0.65; n = 107 No vs. 276 Yes), variants with a downstream EJC showed markedly higher NMD efficiency (median ∼0.72 vs. ∼0.55; P < 0.0001), as did those in highly tolerant genes (pLI < 0.35; n = 1,024 No vs. 2,781 Yes; P < 0.0001) (**panel A**). The same pattern held across allele-frequency classes (**panel B**): ultra-rare variants (n = 931 No vs. 3,234 Yes; P < 0.0001) and rare/common variants (n = 424 No vs. 1,160 Yes; P < 0.0001). CDS-length stratification (**panel C**) produced significant EJC effects in short (<1,289 bp; n = 576 No vs. 1,073 Yes), medium (1,289–2,181 bp; n = 393 No vs. 1,044 Yes), and long (>2,181 bp; n = 232 No vs. 979 Yes) transcripts (all P < 0.0001), and exon-count stratification (**panel D**) yielded significant effects in all four bins (2–5: n = 485 No vs. 410 Yes; 6–10: 359 No vs. 1,009 Yes; 11–20: 272 No vs. 1,114 Yes; >20 exons: 85 No vs. 563 Yes; all P < 0.0001). These observations are consistent with recent studies showing that NMD efficiency differs according to a gene’s tolerance to loss-of-function variants, while PTC position remains a major determinant of NMD outcome^40,41^.

### PTC Position Relative to the Start Codon Shapes NMD Efficiency

We examined how PTC position within the transcript shapes NMD efficiency (**Figure 4**). Across all variants with PTC-to-start distance ≤ 1,000 nt (n = 3,004; **Figure 4A**; Spearman ρ = 0.140), NMD efficiency increased with distance from the translation start site, rising from a median of ∼0.56 within the first 100 nt to a plateau at 200–700 nt. Binning by distance (**Figure 4B**) showed a sharp increase from 0–100 nt (n = 307; median ∼0.56) to 100–200 nt (n = 341; ∼0.67; P < 0.0001), a further rise into 200–300 nt (n = 356; ∼0.74; P < 0.001), a plateau through 300–700 nt (n = 1,252; ns), and a significant decline for PTCs >700 nt (n = 2,683; P < 0.0001)his profile is consistent with potential translation reinitiation downstream of very proximal PTCs, which has been demonstrated mechanistically in multiple disease genes^48–54^ and which would allow short, proximal PTCs to bypass NMD.

**Figure 4.**
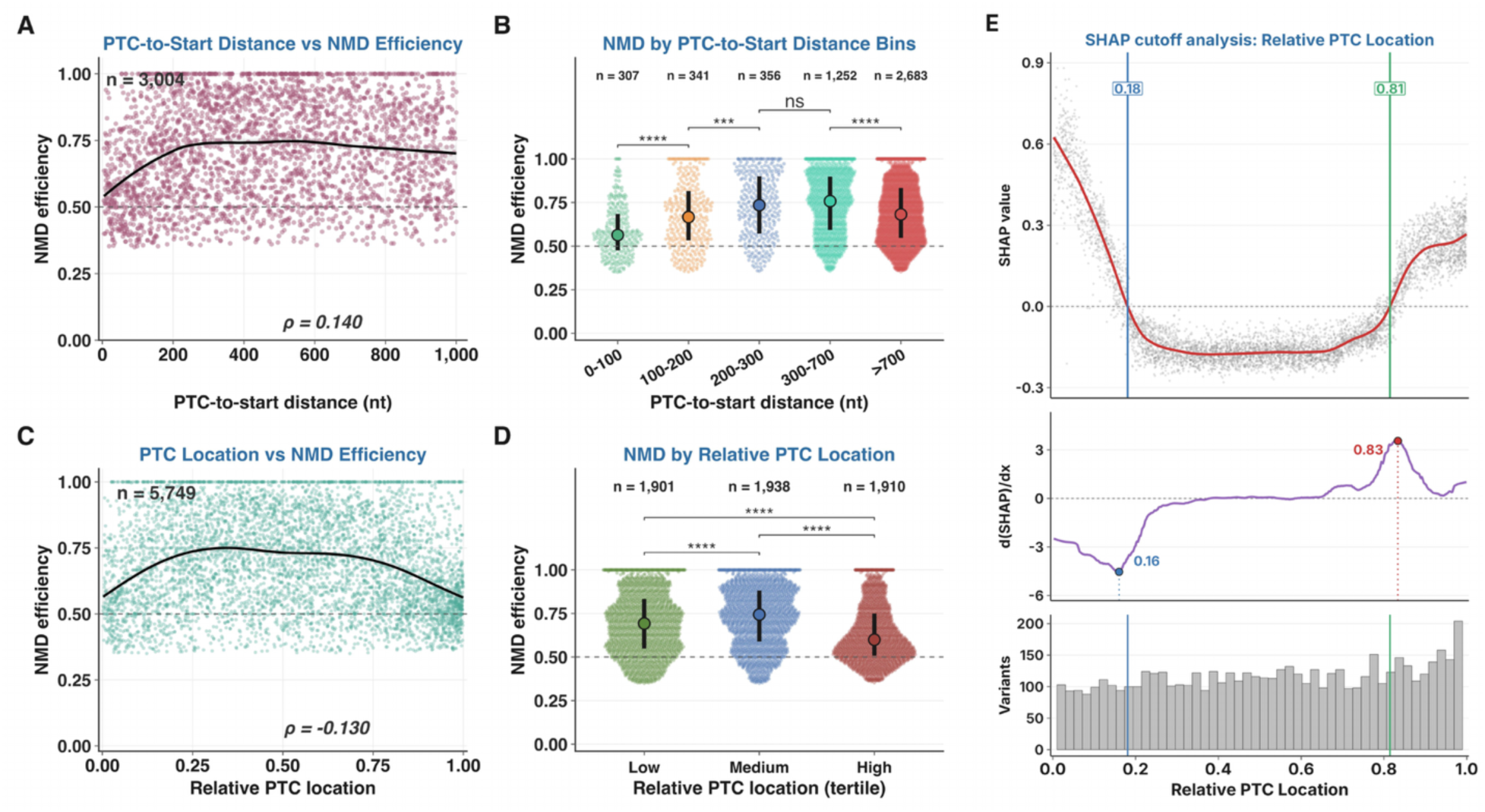
PTC position relative to the start codon and relative PTC location shape NMD efficiency. **(A)** PTC-to-start distance versus NMD efficiency among start-proximal variants (transcripts with ≥ 3 exons and a PTC-bearing exon ≤ 500 bp), shown for distances ≤ 1,000 nt (n = 3,004; Spearman ρ = 0.140); LOESS smoother in black. **(B)** NMD efficiency by PTC-to-start distance bin in the same start-proximal cohort (0– 100 [n = 307], 100–200 [n = 341], 200–300 [n = 356], 300–700 [n = 1,252], >700 nt [n = 2,683]); points are individual variants, large dots mark the median and bars the interquartile range; brackets show comparisons between adjacent bins. **(C)** Relative PTC location (0 = start, 1 = stop) versus NMD efficiency with GAM smoother (n = 5,749; Spearman ρ = −0.130). **(D)** NMD efficiency by relative PTC location tertile (low ≤ 0.375, n = 1,901; medium 0.375–0.714, n = 1,938; high > 0.714, n = 1,910); all three pairwise comparisons **** P < 0.0001. **(E)** SHAP cutoff analysis for relative PTC location from the retrained TrunCat model: positive-to-negative crossover at 0.18 and negative-to-positive crossover at 0.81 (top); rate of change [d(SHAP)/dx], with the steepest decrease at 0.16 and the steepest increase at 0.83 (middle); and the underlying variant distribution (bottom). In-frame downstream AUG and Kozak-context analyses of first-200-bp PTCs are shown in Figure S5. All group comparisons by two-sided Wilcoxon rank-sum test (* P < 0.05, ** P < 0.01, *** P < 0.001, **** P < 0.0001; ns, not significant).

To directly evaluate sequence features associated with the reinitiation hypothesis, we restricted the analysis to variants in the first 200 bp of the coding sequence and asked whether the presence of an in-frame downstream AUG modified NMD efficiency (**Figure S5A**). NMD efficiency did not differ significantly between variants with at least one in-frame downstream AUG (n = 708) and those without (n = 47), indicating that a canonical downstream AUG alone does not predict start-proximal NMD escape at this scale. This finding does not exclude translation reinitiation, which may depend on additional sequence and structural context (proximity, secondary structure, and downstream translational efficiency) documented in disease loci including *BRCA1* and *HBB*^49–53^. Among variants with an in-frame AUG, Kozak-context strength (**Figure S5B**; weak n = 176, moderate n = 231, strong n = 169) also did not significantly differentiate NMD efficiency (all pairwise comparisons not significant). Importantly, these sequence-based analyses identify potential translation-initiation sites but do not directly measure translation reinitiation.

Extending the positional analysis to the full transcript (**Figure 4C**), relative PTC location (0 = start codon, 1 = normal stop) showed a weak but detectable negative correlation with NMD efficiency (n = 5,749; ρ = −0.130), with the generalized additive model (GAM) smoother revealing a modest inverted-U peak around 0.25–0.40 of relative location. In tertile summary (**Figure 4D**; low n = 1,901, medium n = 1,938, high n = 1,910), NMD efficiency differed significantly across tertiles (low vs. medium, P < 0.0001; low vs. high, P < 0.0001; medium vs. high, P < 0.0001), consistent with medium-located PTCs being most efficiently degraded. SHAP cutoff analysis of relative PTC location (**Figure 4E**) confirmed and quantified this inverted-U pattern at the model level: SHAP values were positive (favoring NMD-escape) below a positive-to-negative crossover at 0.18 and above a negative-to-positive crossover at 0.81, and were negative (favoring NMD-sensitive) over the central interval (0.18–0.81) where the model places its strongest sensitivity calls. The rate-of-change panel (middle) shows the steepest descent into the central NMD-sensitive zone near relative location 0.16 and the steepest transition out of it near 0.83, matching the position of the empirical 3ʹ-tertile escape signal in **Figure 4D**. Stratification by CDS length (**Figure S6A, B**) preserved this pattern when relative PTC location was divided into tertiles: PTCs in the 3ʹ tertile had lower NMD efficiency than those in the middle tertile in every CDS length group, whereas the lower efficiency of 5ʹ-tertile PTCs was significant in short and medium transcripts but not in long transcripts. Stratification by exon count (**Figure S6C, D**) showed that relative-location effects are most pronounced in transcripts with 6–20 exons and reduced in very few-exon or very many-exon transcripts, indicating a more complex positional dependence in exon-rich transcripts.

### 3ʹUTR Architecture, Conservation, and RBP Context Are Associated with NMD Efficiency

We evaluated how 3ʹUTR features of original and mutant transcripts (length, composition, intron status, conservation, and RBP-motif content) relate to NMD efficiency (**Figure S7**). 3ʹUTR length stratified for all variants (**S6A**; short log₂ < 10, n = 1,847 vs. long log₂ ≥ 10, n = 1,309) and restricted to last-exon variants (**Figure S7B**; short n = 504 vs. long n = 371) showed only modest within-stratum differences (both ns), consistent with a broad rather than cliff-edge contribution of 3ʹUTR length to NMD efficiency. 3ʹUTR composition (**Figure S7C**, D) showed a significant directional effect: higher AU content (**panel C**) was associated with increased NMD efficiency (low vs. high, P < 0.0001), while higher UC content (**panel D**) showed modest non-monotonic associations (low vs. medium P < 0.05; medium vs. high ns). Among last-exon variants, transcripts with a 3ʹUTR intron (n = 22) showed higher NMD efficiency than those without one (n = 891; median ≈ 0.63 vs. ≈ 0.54; Wilcoxon P < 0.01; **Figure S7E**). This is aligned with a prior study^94^ that an exon junction downstream of the stop codon may lead to increased NMD efficiency, although the 3ʹUTR-intron group is small.

3ʹUTR conservation had a clear directional effect (**Figure S7F**): variants in transcripts with highly conserved first-200-nt new 3ʹUTRs (PhastCons ≥ 0.5; n = 2,325) showed significantly higher mean NMD efficiency (∼0.708) than those with low conservation (<0.1; n = 742; ∼0.638; P < 0.0001), with intermediate conservation (0.1–0.5; n = 147; ∼0.659) showing a smaller medium-to-high difference (P < 0.001). RBP motif analysis within the 3ʹUTR (**Figure S7G**) identified five Bonferroni-significant motifs (SAMD4A, CELF6, HNRNPH2, HNRNPH1, and HNRNPH3), all with negative Δ (“inhibits NMD”).

### 5ʹUTR Features Are Weak Determinants of NMD Efficiency

In contrast to the 3ʹUTR, 5ʹUTR features contributed little to NMD outcomes (**Figure S8**). 5ʹUTR AU-content tertiles (**Figure S8A**; n ≈ 1,055 per tertile) showed no significant associations, and 5ʹUTR UC content (**Figure S8B**) produced only a small shift (medium vs. high P < 0.05). The presence of a 5ʹUTR uORF (**Figure S8C**; n = 1,116 without vs. 2,051 with) did not significantly alter NMD efficiency, whereas transcripts with a 5ʹUTR intron (**Figure S8D**; n = 1,058 with vs. 2,156 without) showed slightly lower NMD efficiency (P < 0.001). 5ʹUTR PhastCons conservation (**Figure S8E**) showed only a subtle pattern. Together, these results indicate that 5ʹUTR properties except 5ʹUTR intron and 5ʹUTR PhastCons conservation are not major determinants of NMD efficiency in germline PTC-variants.

### Global Transcript Architecture, mRNA Stability, and Expression Further Refine NMD Prediction

We evaluated a broader panel of transcript-level features (CDS length, exon count, expression level, mRNA half-life, isoform diversity, and CDS composition) to characterize their marginal effects on NMD efficiency (**Figure 5**; one observation per transcript, n = 3,214). CDS length showed no significant association with NMD efficiency at the transcript level (**Figure S3C**). The directional behavior of CDS length, which appeared among the top 20 features (log2_CDS, mean |SHAP| = 0.036), is dissected in **Figure S9A**: per-variant SHAP dependence is weakly negative for log₂ CDS < 11 and rises above a bootstrap zero-crossing at log₂ CDS = 11.02 (95% CI 10.98– 11.05; CDS ≈ 2.1 kb; **Figure S9D**), pushing predictions toward NMD escape in long transcripts. The effect is driven by the upstream-EJC (NMD-sensitive) subset (crossing at 11.01; **Figure S9G**), with no robust crossing in last-exon or penultimate-last-50 bp variants, indicating that long CDS length contributes to predicted NMD escape specifically among variants with a downstream EJC. Exon count (**Figure 5A**) showed a clear effect: transcripts in the lowest exon-count tertile (2–7 exons; T1; n = 1,127) had significantly lower median NMD efficiency (∼0.63) than those in the middle tertile (8–13 exons; T2; n = 1,017; ∼0.71; P < 0.0001), after which the distribution plateaued (T2 vs. T3, ns). Median transcript expression (**Figure 5B**) did not differ between the lowest and middle tertiles but showed a subtle, significant increase in the highest tertile (median ∼0.67 in T2 vs. ∼0.70 in T3; P < 0.01).

**Figure 5.**
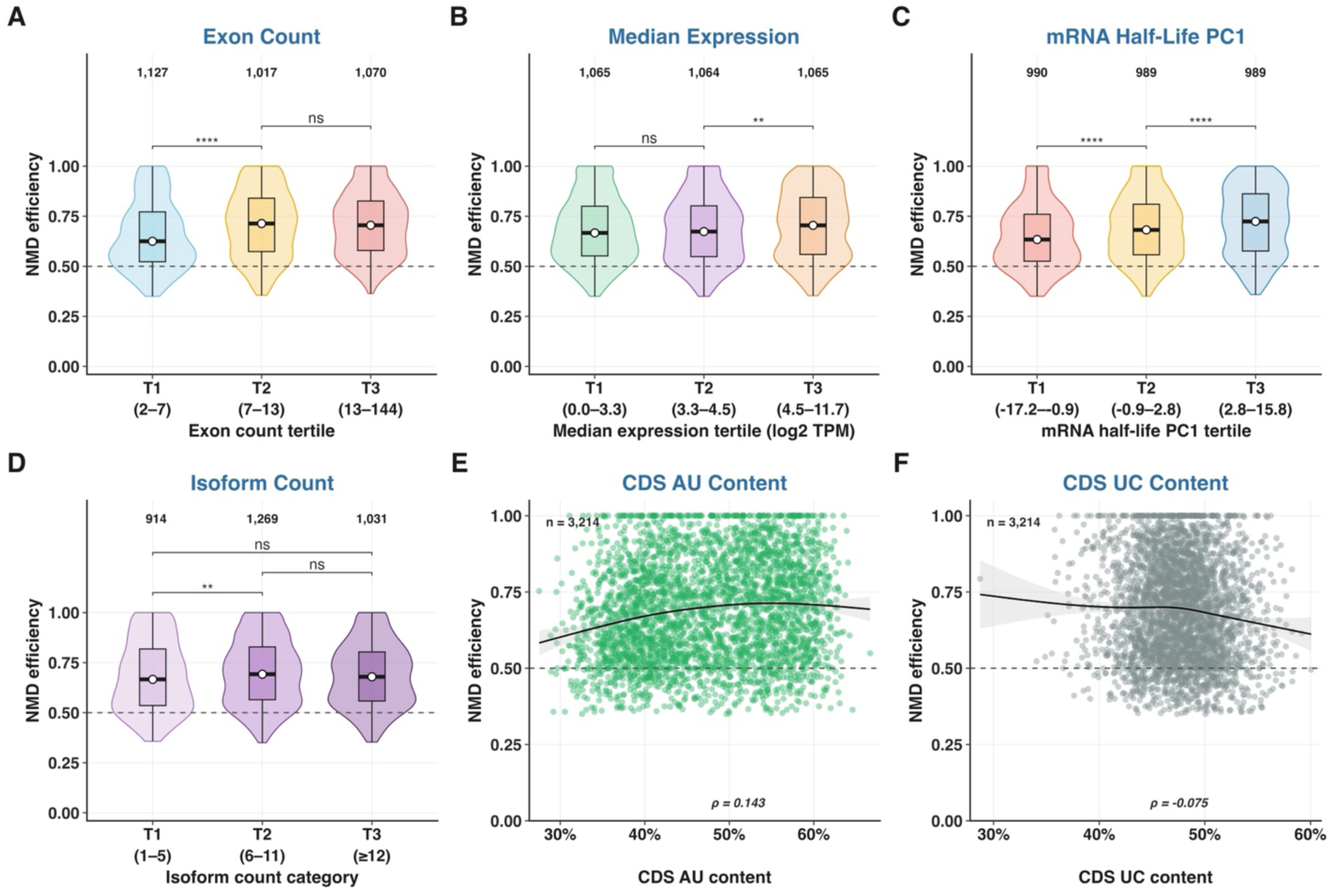
Transcript architecture, mRNA stability, expression, and CDS composition are associated with NMD efficiency. All analyses are transcript-level: each transcript contributes one observation, the median NMD efficiency across its variants (n = 3,214 transcripts), and all bins are defined across transcripts. **(A)** NMD efficiency by exon count tertile (T1, 2–7 exons, n = 1,127; T2, 8–13, n = 1,017; T3, 14–144, n = 1,070). **(B)** NMD efficiency by median transcript expression tertile (log2 TPM; n = 1,065, 1,064, and 1,065). **(C)** NMD efficiency by mRNA half-life PC1 tertile (n = 990, 989, and 989). **(D)** NMD efficiency by isoform count category (1–5, n = 914; 6–11, n = 1,269; ≥12, n = 1,031). **(E, F)** Continuous relationship between CDS nucleotide composition, AU content (E) and UC content (F), and NMD efficiency, with GAM smoother and Spearman ρ (E, ρ = 0.143; F, ρ = −0.075; n = 3,214). Violins show the distribution, boxes the interquartile range, and white circles the median. Group comparisons in A–D use two-sided Wilcoxon rank-sum tests between adjacent bins (A–C: T1–T2, T2–T3; D: 1–5 vs 6–11, 6–11 vs ≥12, and 1–5 vs ≥12) (* P < 0.05, ** P < 0.01, *** P < 0.001, **** P < 0.0001; ns, not significant).

mRNA half-life (PC1)^89^ (**Figure 5C**) produced a striking monotonic effect: transcripts in the lowest half-life PC1 tertile (T1, shortest half-lives) showed significantly lower NMD efficiency (median ∼0.63) than those in the highest tertile (T3, longest half-lives; ∼0.72), with both adjacent-tertile steps significant (T1→T2 and T2→T3, both P < 0.0001). This is consistent with mRNA half-life being the third most important feature in TrunCat (mean |SHAP| = 0.160; **Figure 2D**): transcripts with longer intrinsic half-lives may undergo more rounds of translation and therefore have more opportunities for NMD engagement, while very unstable transcripts are rapidly degraded by alternative pathways that outpace NMD. Isoform count (**Figure 5D**) showed a modest effect at the low end (1–5 vs. 6–11 isoforms, P < 0.01), with no further difference at ≥ 12 isoforms, consistent with established coupling of alternative splicing to NMD (AS-NMD) in which highly alternatively spliced genes generate more NMD-target isoforms^95,96^. SHAP analyses of two of the strongest non-canonical contributors are provided in **Figures S10** and **S11**. For median transcript expression (mean |SHAP| = 0.049; **Figure S10**), per-variant SHAP values shift from negative (NMD-triggering) at low expression to positive/neutral (NMD-escape) at high expression. mRNA half-life PC1 (mean |SHAP| = 0.160; **Figure S11**) showed an even more pronounced monotonic dependence: SHAP values are strongly positive (favoring NMD-escape) at low PC1 and shift to strongly negative (favoring NMD-sensitive) at high PC1. Together, these analyses establish that CDS length, median expression, and mRNA half-life act as graded, EJC-class-modulated determinants of NMD efficiency at the model level.

The continuous relationship between CDS nucleotide composition and NMD efficiency (**Figure 5E, F**; n = 3,214 transcripts) showed a weak but significant positive correlation with CDS AU content (ρ = 0.143, P = 3.0 × 10⁻^1^⁶; **Figure 5E**) with a mild inverted-U peaking near 50–55% AU, and the mirror-image weak negative correlation with CDS UC content (ρ = −0.075, P = 1.9 × 10⁻⁵; **Figure 5F**).

### PTC-to-EJC Distance, Local Conservation, and RBP Motifs Provide Additional Signals of NMD Efficiency

To dissect the local sequence environment that determines NMD outcome, we examined PTC-to-EJC distance, sequence conservation in the PTC-to-EJC interval, and RBP-motif content within three windows (PTC-to-EJC, PTC ± 100 nt, and EJC ± 100 nt) (**Figure 6**). Across all variants with a downstream EJC and not in the first 200 bp of a transcript (n = 3,684), PTC-to-EJC distance showed a weak overall negative association with NMD efficiency at the global scale (**Figure 6A**; Spearman ρ = −0.073, P = 9.1×10⁻⁶), but the LOESS smoother revealed a clear non-monotonic structure rather than a flat profile: mean efficiency was highest (∼0.78) in the proximal range (0-250 nt), declined visibly across the intermediate range (∼250–750 nt) to a minimum near ∼0.65, and partially recovered/plateaued beyond ∼750 nt.

**Figure 6.**
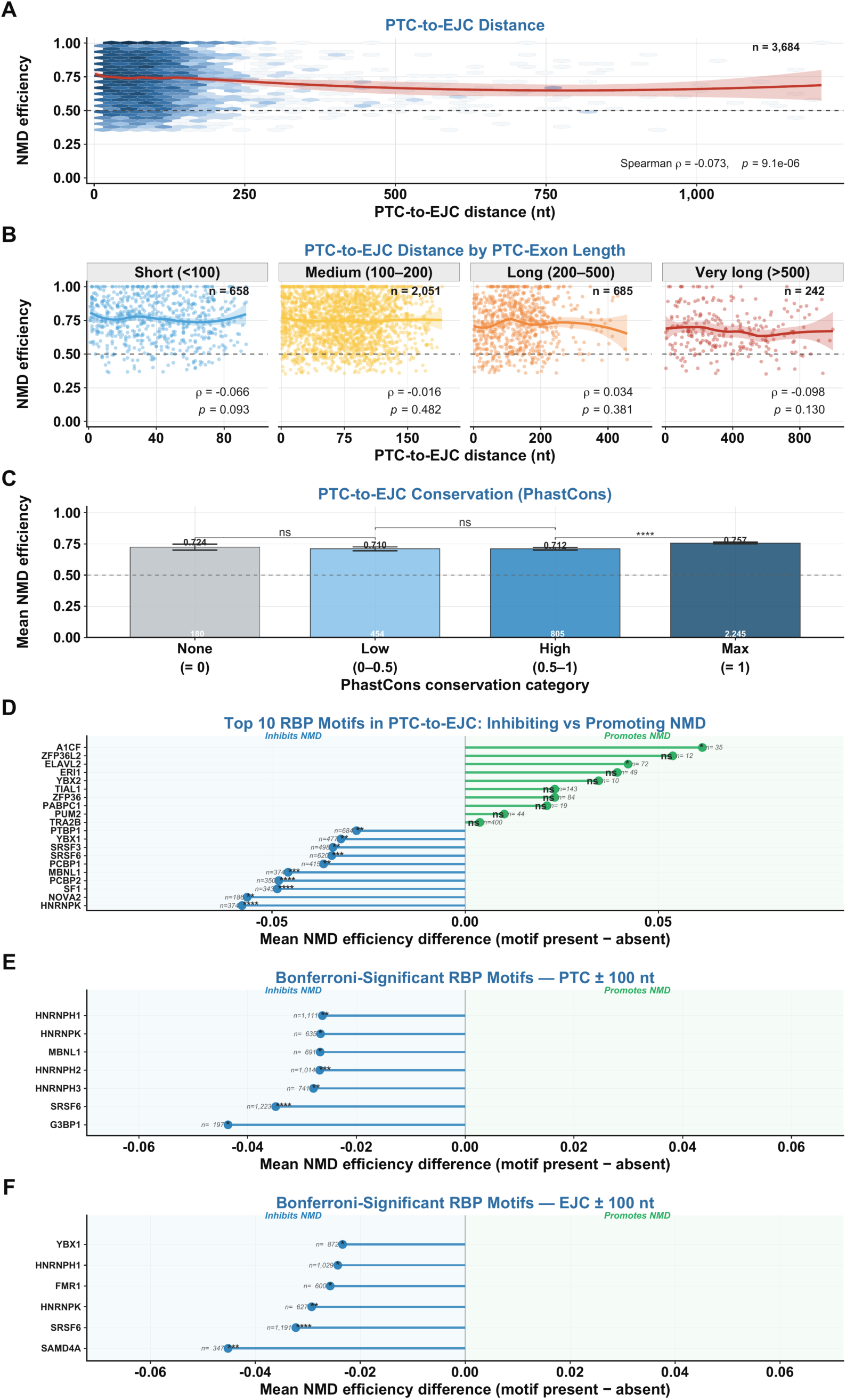
Local PTC sequence context: PTC-to-EJC distance, exon-length stratification, conservation, and RBP-motif environment. All panels use the variant-level subset of nonsense variants with a downstream EJC (PTC upstream of the last EJC) and the PTC located outside the first 200 bp of the CDS (n = 3,684). **(A)** PTC-to-EJC distance versus NMD efficiency for all such variants; LOESS smoother (red) with 95% CI (Spearman ρ = −0.073, P = 9.1×10⁻⁶). **(B)** The same relationship stratified by PTC-host-exon length: short (<100 bp; n = 658; ρ = −0.066, P = 0.093), medium (100–200 bp; n = 2,051; ρ = −0.016, P = 0.482), long (200–500 bp; n = 685; ρ = +0.034, P = 0.381), and very long (>500 bp; n = 242; ρ = −0.098, P = 0.130); each Spearman ρ is computed within the facet’s displayed distance window. **(C)** Mean NMD efficiency (± 95% CI) by PhastCons conservation of the PTC-to-EJC interval (None = 0, n = 180; Low 0– 0.5, n = 454; High 0.5–1, n = 805; Max = 1, n = 2,245); among adjacent-category Wilcoxon comparisons, only the High→Max step is significant (**** P < 0.0001), while None→Low and Low→High are not significant (ns). **(D)** Top 10 inhibiting and top 10 promoting RBP motifs in the PTC-to-EJC window, selected by significance and displayed in order of mean NMD-efficiency difference (motif present − absent); inhibitors (blue) are all Bonferroni-significant, whereas none of the promoters (green) is Bonferroni-significant (promoters are labeled with unadjusted significance). **(E)** All Bonferroni-significant RBP motifs in the PTC ± 100 nt window (7 inhibitors; no significant promoters). **(F)** All Bonferroni-significant RBP motifs in the EJC ± 100 nt window (6 inhibitors; no significant promoters). RBP-motif comparisons (D–F) use two-sided Wilcoxon rank-sum tests with Bonferroni correction. Significance: * P < 0.05, ** P < 0.01, *** P < 0.001, **** P < 0.0001; ns, not significant.

Stratifying by PTC-host-exon length confirmed and refined this interpretation (**Figure 6B**). Although the per-stratum Spearman correlations were modest, the within-stratum LOESS smoothers consistently showed an early plateau followed by a visible decrease in NMD efficiency at the longer end of each accessible distance range, mirroring the global pattern in **Figure 6A**. The decline was most readily apparent in medium (100–200 bp), long (200–500 bp), and very long (>500 bp) host exons, where larger PTC-to-EJC distances are achievable.

Sequence conservation of the PTC-to-EJC interval retained a clear effect (**Figure 6C**). Variants with no conserved positions (PhastCons = 0; n = 180) had mean NMD efficiency of 0.724, low-conservation intervals (0–0.5; n = 454) 0.710, high-conservation intervals (0.5–1; n = 805) 0.712, and maximally conserved intervals (=1; n = 2,245) 0.757. The first three categories were statistically indistinguishable, while maximally conserved intervals showed a small but highly significant elevation over the other categories (High vs. Max P < 0.0001). Related conservation features around the downstream EJC and in the PTC-to-EJC interval (phastcons_ejc_100bp_median, phylop_ejc_100bp_median, and phylop_ptc_to_ejc_median) were among the top-20 contributors to TrunCat (**Figure 2D**). RBP-motif analyses revealed a rich and reproducible regulatory landscape across all three windows (**Figure 6D–F**). In the PTC-to-EJC window (**Figure 6D**), the panel shows the ten strongest inhibiting and ten strongest promoting RBP motifs, selected by significance and displayed by mean NMD-efficiency difference. The inhibitors were dominated by hnRNP-family and related splicing and RNA-processing factors (negative Δ; 19 motifs were Bonferroni-significant by Wilcoxon rank-sum test, including all ten displayed inhibitors): the top five by effect size were HNRNPK (n = 374), NOVA2 (n = 186), SF1 (n = 343), PCBP2 (n = 350), and MBNL1 (n = 374). In the PTC ± 100 nt window (**Figure 6E**), seven inhibitors reached Bonferroni significance, with G3BP1 (n = 197), SRSF6 (n = 1,223), HNRNPH3 (n = 741), HNRNPH2 (n = 1,014), and MBNL1 (n = 691) the top five by effect size, and no motif was a Bonferroni-significant promoter. In the EJC ± 100 nt window (**Figure 6F**), the picture was likewise uniformly inhibitory: six motifs (SAMD4A, SRSF6, HNRNPK, FMR1, HNRNPH1, and YBX1) were significantly associated with reduced NMD efficiency, and no motif emerged as a Bonferroni-significant promoter. The recurrent recovery of hnRNP-family motifs (HNRNPK, HNRNPH1, HNRNPH2, and HNRNPH3) together with SRSF6, MBNL1, and YBX1 as inhibitors across these windows suggests coordinated cis-regulatory families that modulate NMD independently of canonical EJC positioning, consistent with biochemical and tissue-resolved data demonstrating that hnRNP proteins and stress-granule–associated factors act as cell-context-dependent modifiers of UPF1 deposition and NMD-substrate selection^58,91,97–100^.

### Predicted NMD-Escape PTCs Are Enriched among ClinVar VUS and Rare-Disease Variants

To explore population- and clinical-scale applications of TrunCat, we generated per-variant predictions on four external variant sets that lack ASE-derived NMD labels: gnomAD (n = 137,857; 38.4% predicted escape), ClinVar pathogenic/likely-pathogenic nonsense variants (P/LP; n = 23,856; 28.4% predicted escape), ClinVar variants of uncertain significance (VUS; n = 3,445; 48.3% predicted escape), and a rare-disease cohort from the GREGoR Consortium (n=2,548; 45.5% predicted escape) (**Figure 7**). Predicted escape-probability distributions were broadly bimodal across all four datasets (**Figure 7A**), with clear separation around the decision threshold τ = 0.47. ClinVar P/LP predictions were shifted toward the NMD-sensitive mode relative to gnomAD, consistent with the enrichment of canonical loss-of-function (NMD-sensitive) alleles among high-confidence P/LP variants. Strikingly, ClinVar VUS predictions showed a distinct distribution shifted toward higher escape probabilities (**Figure 7B**: 51.7% NMD-sensitive and 48.3% escape), yielding an escape-class composition ∼1.7× that of P/LP variants (48.3% vs. 28.4%). The GREGoR rare-disease cohort showed an intermediate but elevated escape fraction (45.5% escape, 54.5% NMD-sensitive), closer to the VUS distribution than to either gnomAD or P/LP, consistent with enrichment of NMD-escape alleles in clinically ascertained probands whose causal variant has not yet been definitively classified.

**Figure 7.**
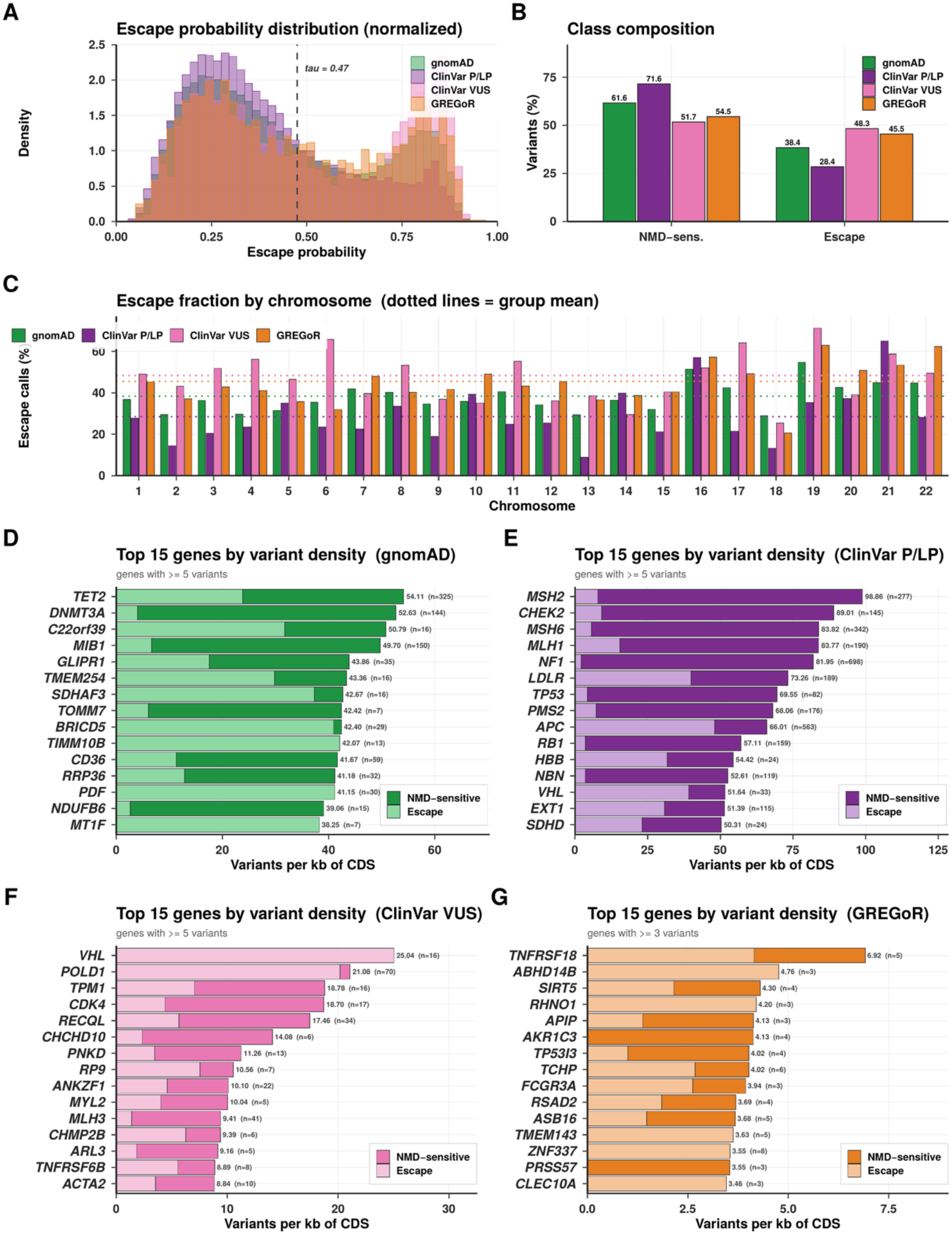
Population- and clinical-scale NMD-escape prediction across gnomAD, ClinVar P/LP, ClinVar VUS, and the GREGoR rare-disease cohort. (A) Normalized escape-probability distributions for the four datasets; dashed line marks the decision threshold τ = 0.47. (B) Class composition (% NMD-sensitive vs. % escape) in each dataset: gnomAD 61.6/38.4, ClinVar P/LP 71.6/28.4, ClinVar VUS 51.7/48.3, GREGoR 54.5/45.5. (C) Per-chromosome predicted escape fractions; dotted lines indicate dataset means. (D–G) Top 15 genes by variant density (variants per kilobase of coding sequence, among genes with ≥ 5 variants, or ≥ 3 for GREGoR) in gnomAD (D), ClinVar P/LP (E), ClinVar VUS (F), and GREGoR (G), with each bar split into predicted NMD-sensitive (dark shading) versus NMD-escape (light shading) calls and the total density and variant count annotated to the right of each bar.

Predicted escape fractions varied by chromosome (**Figure 7C**) but stayed close to dataset means for most autosomes, with modest elevation on chromosomes 16, 19, 21, and 22, likely reflecting gene-content composition. Across datasets, ClinVar VUS and the GREGoR rare-disease cohort showed consistently higher per-chromosome escape fractions than P/LP variants, reinforcing the overall shift toward escape seen in the class composition (**Figure 7B**) and supporting the hypothesis that NMD-escape PTCs are over-represented among variants of uncertain or unresolved clinical significance.

Top genes by germline nonsense variant density (**Figure 7D–G**) highlighted disease-relevant loci with distinct sensitivity/escape profiles across the four datasets. In gnomAD (**Figure 7D**), the top five genes by variant density were *TET2*, *DNMT3A*, *C22orf39*, *MIB1*, and *GLIPR1*. In ClinVar P/LP (**Figure 7E**), the top five genes by variant density were *MSH2*, *CHEK2*, *MSH6*, *MLH1*, and *NF1*, all classical tumor-suppressor or mechanistically characterized loss-of-function genes. In ClinVar VUS (**Figure 7F**), the leading genes were *VHL*, *POLD1*, *TPM1*, *CDK4*, and *RECQL*. The GREGoR rare-disease cohort (**Figure 7G**) showed a notably different gene spectrum with lower variant densities, led by *TNFRSF18*, *ABHD14B*, *SIRT5*, *RHNO1*, and *APIP*, consistent with the rare disease ascertainment of this cohort and the enrichment of novel or under-characterized loci within it. Across all four panels, NMD-sensitive (dark) and NMD-escape (light) calls were heterogeneously distributed within individual genes, illustrating that even classical haploinsufficient disease genes harbor a mixture of canonical loss-of-function and predicted-escape alleles.

Taken together, these results demonstrate that TrunCat produces calibrated, interpretable predictions at population scale across both gnomAD, P/LP and VUS variants (ClinVar P/LP and ClinVar VUS) clinically ascertained rare-disease (GREGoR) variant sets, and identifies a substantial fraction of ClinVar VUS and GREGoR variants as predicted NMD-escape alleles, providing a candidate set for mechanistic follow-up to resolve their pathogenicity^41^. To assess clinical actionability in the most disease-relevant subset of variants, we further restricted the same four-dataset comparison to autosomal-dominant, highly loss-of-function intolerant genes (LOEUF < 0.6, pLI ≥ 0.9; 778 genes); the resulting escape-probability landscape and per-chromosome and per-gene breakdowns are provided in **Figure S12**. Within this constrained-gene subset, ClinVar VUS again showed the highest predicted escape fraction (43.4%, versus 33.8% for P/LP and 32.7% for gnomAD), whereas the GREGoR cohort showed a lower escape fraction (30.2%) than in the full gene set.

### TrunKitten Predictions Are Concordant with Experimentally Measured NMD Efficiency in LMNA

To evaluate TrunKitten against experimentally measured NMD activity, we applied the eight-feature model, without retraining, to the *LMNA* saturation genome editing (SGE) dataset of Cortázar et al.^92^, in which PTCs were introduced across the first, internal, and penultimate exons of *LMNA* and NMD efficiency was quantified with and without pharmacological NMD inhibition. Of the 86 engineered PTCs that can arise from the endogenous codon by a single nucleotide substitution, 84 passed the experimental variability filter. TrunKitten-predicted NMD probability correlated positively with experimentally measured NMD efficiency (Spearman ρ = 0.500, P = 1.3 × 10⁻⁶; **Figure S13A**). When variants were classified at the TrunKitten decision threshold, those predicted to trigger NMD (n = 37; exons 4, 7, and 10) showed higher experimental NMD efficiency than those predicted to escape NMD (n = 47; exons 1, 10, and 11; median 0.86 vs. 0.75; Wilcoxon rank-sum P = 4.9 × 10⁻⁵; **Figure S13B**).

## Discussion

Across 5,749 high-quality heterozygous nonsense variants in 3,214 different transcripts in 10,306 TOPMed individual samples, the canonical EJC-dependent rule^17,19^ emerged as the dominant predictor of NMD efficiency (**Figure 3A–C**; “EJC downstream of PTC” mean |SHAP| = 0.515), and its dominance held across every stratum of gene-level constraint, allele frequency, CDS length, and exon count (**Figure S4**). Layered on top of this binary rule, our SHAP-ranked top-20 features identify a coherent set of quantitative refinements that together raise out-of-fold ROC-AUC to 0.777 (PR-AUC 0.733) (**Figure 2F**). The five most informative non-canonical contributors were (i) relative PTC location within the transcript (mean |SHAP| = 0.193), which produced an inverted-U dependence with maximum NMD efficiency in the central tertile (**Figure 4C–D**) and reflects the combination of 5ʹ-proximal escape via potential translation reinitiation^49–54^ and 3ʹ-proximal escape within the EJC-free last-exon window^22,37,38^; (ii) mRNA half-life PC1^93^ (mean |SHAP| = 0.160), which showed a monotonic positive association with NMD efficiency (**Figure 5C**), consistent with the established coupling between NMD and translation^21^; (iii) CDS AU content (mean |SHAP| = 0.153), with AU-rich transcripts being preferred substrates; (iv) PTC-to-EJC distance, which showed a graded, nonlinear relationship with NMD efficiency, with a local minimum in the fitted curve at ≈18 nt (**Figure 3D**), consistent with the canonical 50–55-nt boundary reflecting a continuous distance-dependent relationship rather than a strict binary threshold; and (v) sequence conservation in the PTC-to-EJC interval and around the 3ʹUTR, with “phastcons_new3utr_first200_median”, “phastcons_ejc_100bp_median”, “phylop_ejc_100bp_median”, and “phylop_ptc_to_ejc_median” all in the top-20 (**Figure 2D**). Although translation reinitiation has been experimentally shown to protect transcripts carrying start-proximal PTCs from NMD, including reinitiation at downstream AUG codons^49,51^, neither the presence of an in-frame downstream AUG nor its Kozak-context strength was significantly associated with NMD efficiency in our population-scale analysis (**Figure S5A–B**). These findings do not exclude translation reinitiation as a mechanism of start-proximal NMD escape; rather, they indicate that the canonical downstream-AUG sequence features examined here are insufficient on their own to explain the observed escape pattern. Our sequence-based analysis identifies potential initiation sites but does not directly measure ribosomal reinitiation. Furthermore, mammalian translation initiation is not restricted to AUG codons, and non-AUG initiation is strongly influenced by local sequence and structural context^101^, raising the possibility that non-AUG reinitiation events not captured by our AUG-based analysis may also contribute to start-proximal escape. A complementary mechanism, not mutually exclusive with reinitiation, is the closed-loop model proposed by Silva et al.^47^, in which a 5ʹ–3ʹ closed-loop mRNA conformation positions the cytoplasmic poly(A)-binding protein (PABPC1) in proximity to start-proximal PTCs, where it antagonizes the NMD machinery and inhibits decay; saturation genome editing data from our group are consistent with this PABPC1-proximity model as an explanation for NMD inhibition at the 5ʹ ends of transcripts^92^. RBP-motif analyses (**Figure 6D–F**) further recovered a coordinated cis-regulatory landscape in which hnRNP-family motifs (HNRNPK, HNRNPH1/2/3) together with SRSF6, MBNL1, YBX1, and G3BP1 recurrently inhibit NMD across the PTC-to-EJC, PTC ± 100 nt, and EJC ± 100 nt windows, consistent with biochemical evidence that hnRNP-family factors (including HNRNPL binding to GC-rich 3ʹUTR elements) and stress-granule–associated factors act as cell-context-dependent modifiers of UPF1 deposition^58,91,97–100^.

The graded PTC-to-EJC relationship suggests that the canonical 50–55-nt rule represents a strong positional tendency rather than an absolute molecular boundary. The fitted relationship reached a local minimum at approximately 18 nt before NMD efficiency increased with greater PTC-to-EJC distance. We interpret this short-distance feature as an empirical characteristic of the population-level distance-response relationship rather than a new discrete threshold for NMD. This interpretation is consistent with Cortázar et al.^92^, who similarly observed a quantitative rather than strictly binary relationship between PTC position and NMD activity. Gangras et al.^94^ further demonstrated that a subset of transcripts containing 3′UTR introns within 50 nt of the normal termination codon can undergo EJC-dependent NMD, indicating that downstream exon junctions can promote NMD in contexts not fully captured by a strict 50-nt boundary.

Motivated by these findings, we examined whether 3′UTR introns also influence NMD of PTCs in the last coding exon, which are generally expected to escape NMD because no exon junction lies downstream of the PTC. Among last-exon variants, transcripts containing a 3′UTR intron (n = 22) showed higher NMD efficiency than those without one (n = 891; median ≈ 0.63 vs. ≈ 0.54; Wilcoxon P < 0.01; **Figure S7E**). This is consistent with a 3′UTR intron placing an exon junction downstream of the PTC and thereby restoring EJC-dependent NMD, although this result should be interpreted cautiously given the small number of 3′UTR-intron-containing transcripts. Together, these observations are consistent with a model in which the position of downstream exon junctions, including those in the 3′UTR, contributes quantitatively to NMD efficiency, while transcript context may further modify this relationship. Further studies with larger numbers of 3′UTR-intron-containing transcripts will be required to determine the underlying mechanism.

The association between mRNA half-life and NMD efficiency should be considered in the broader context of the coupling between NMD and translation. Early studies provided evidence that mammalian NMD occurs during a pioneer round of translation on transcripts associated with the nuclear cap-binding complex (CBC)^21^. However, subsequent studies demonstrated that NMD is not strictly restricted to this stage. Durand and Lykke-Andersen demonstrated that NMD substrates remain susceptible to decay during eIF4F-dependent translation, including after exchange of the CBC for eIF4F, indicating that NMD can act on mRNAs beyond the pioneer round^102^. Furthermore, using single-molecule imaging to directly monitor translation and decay of individual mRNAs, Hoek et al. demonstrated that NMD can occur during multiple rounds of translation, with each terminating ribosome having a similar probability of inducing NMD^103^. Together, these findings establish that NMD is coupled to translation but is not strictly confined to the pioneer round. Thus, the association between mRNA half-life and NMD efficiency observed here is consistent with the translation-dependent nature of NMD but does not imply a strict requirement for pioneer-round translation.

These findings recapitulate and extend prior landmark analyses. Lindeboom et al. (2016, 2019) used somatic variants in 9,769 tumors to derive an integrated multi-feature NMD model and the rule-based NMDetective resource^37,38^, establishing non-canonical determinants, first-150-nt escape, long-middle-exon escape, last-EJC proximity, and a modulating role for mRNA half-life, that we recover in their entirety. Three contemporaneous studies define the current state of the art: Iha et al. integrated multi-nucleotide variant rescue, ribosome occupancy, and isoform expression to improve rule-based annotation by 12.0% on 1,086 individuals^29^; Saadat and Fellay developed NMDEP, an integrative model combining sequence embeddings with curated biological features^70^; and Veiner, Toledano and colleagues developed NMDetective-AI by fine-tuning the Orthrus mRNA foundation model on ∼14,000 somatic PTCs and validating on ∼11,000 germline GTEx PTCs (Spearman ρ ∼ 0.56–0.67), coupled to three deep mutational scanning libraries^31^. TrunCat and NMDetective-AI^31^ converge on the same biological picture, the 50-nt rule is a graded logistic transition, long exons partially evade NMD, and start-proximal escape is real and reinitiation-mediated, but they occupy complementary niches: NMDetective-AI^31^ maximizes expressive capacity through learned sequence embeddings and is well suited to in silico saturation mutagenesis, whereas TrunCat operates on 619 hand-engineered features, returns calibrated SHAP attributions for every prediction, and is trained exclusively on germline ASE-derived NMD efficiency, the regime most directly relevant to ACMG/AMP PVS1 classification^32,33^. Locus-saturation experiments by Cortázar et al. (2025)^92^, who measured NMD activity for 722 PTCs across five exons of *LMNA*^92^, independently validate three of our population-scale findings: a quantitative rather than binary PTC-to-EJC distance dependence, near-absent NMD in the first ∼21 codons (mirroring our 0–100 vs. 100–200 nt PTC-to-start transition; **Figure 4A–B**), and readthrough-permissive contexts (the TGA-CT motif) that are represented in TrunCat by its HEK293T readthrough score feature.

Applied at scale, TrunCat predicts that 48.3% of 3,445 ClinVar VUS carry NMD-escape PTCs, compared with only 28.4% of 23,856 ClinVar pathogenic/likely-pathogenic variants and 38.4% of 137,857 gnomAD variants (**Figure 7**), a 1.7-fold VUS-vs-P/LP enrichment that nominates a candidate set for mechanistic follow-up and potential reclassification. Application to a clinically ascertained rare-disease cohort from the GREGoR Consortium yielded an intermediate but elevated escape fraction (45.5%), closer to the VUS distribution than to either gnomAD or P/LP and consistent with enrichment of NMD-escape alleles among unsolved or partially solved rare-disease probands. Restricting these comparisons to autosomal-dominant, highly loss-of-function intolerant genes (LOEUF < 0.6, pLI ≥ 0.9; 778 genes; **Figure S12**) kept ClinVar VUS as the dataset with the highest predicted escape fraction and identified a tractable, disease-relevant subset of predicted NMD-escape alleles for prioritization. Because NMD has been described as both “friend and foe” in genetic disease, alleviating phenotypes by clearing harmful dominant-negative or gain-of-function truncated proteins, or aggravating phenotypes by eliminating partially functional transcripts that could have provided residual gene activity^45,58–62,95^, calibrated per-variant escape probabilities allow stratification of patients into those whose disease is likely to be NMD-aggravated (and therefore expected to benefit from NMD inhibition or translational readthrough^32,59,62–67^) versus NMD-ameliorated, where NMD inhibition would risk producing harmful truncated proteins. The recent genome-scale readthrough resource of Toledano et al. across ∼5,800 pathogenic stop codons^64^ complements TrunCat’s NMD predictions for selecting PTC alleles most likely to benefit from small-molecule readthrough therapy.

Several limitations should be acknowledged. First, TrunCat is trained on whole-blood RNA-seq, and recent population- and tissue-resolved analyses have shown that NMD efficiency varies systematically between tissues and individuals, with reproductive and nervous-system tissues showing lower baseline activity than digestive tissues^57,58^; predictions for variants whose disease relevance is concentrated in low-NMD tissues may therefore be biased. Second, ASE is an indirect measure of NMD efficiency that cannot fully separate NMD from other mechanisms of allelic imbalance. Third, purifying selection in the TOPMed cohort biases the variant distribution toward less deleterious PTCs, potentially under-sampling the high-efficiency end of the spectrum. Future work integrating tissue-matched RNA-seq (e.g., GTEx), allele-specific nascent-transcription data, and long-read sequencing will refine prediction further, while expanded training in patient-derived cohorts will strengthen clinical portability. With these caveats, TrunCat and its accompanying ClinVar VUS predictions provide a scalable, mechanistically grounded resource for PTC-variant interpretation that complements rule-based tools and is immediately usable within ACMG/AMP workflows.

Identifying novel NMD prediction rules and developing a more accurate NMD prediction model better informs genetic and clinical research, potentially leading to more precise diagnosis and treatment strategies for diseases caused by PTC-variants. Leveraging large-scale genomic data such as TOPMed provides deeper understanding of RNA biology and novel insights into NMD outcomes and mechanisms. Our integrated framework, combining rigorous feature engineering with interpretable gradient boosting and orthogonally validated by saturation-genome-editing measurements^92^ and contemporaneous deep mutational scanning studies^31^ offers a scalable approach to NMD outcome prediction that complements and extends the canonical EJC-based rule and can be directly applied to population- and clinical-scale variant sets, including ClinVar VUS.

## Data and code availability

All code and annotations, including data processing, feature engineering, and model training, are available at https://github.com/CobanAkdemirlab/NMDpredictionmodel. TOPMed data are available through dbGaP (accession phs001024). gnomAD data are publicly available at https://gnomad.broadinstitute.org/. ClinVar data are available at https://www.ncbi.nlm.nih.gov/clinvar/. GREGoR data are available through dbGaP per consortium data access policies. Further information and requests for resources should be directed to and will be fulfilled by the lead contact, Zeynep Coban-Akdemir with her e-mail address.

## Acknowledgments

This work was supported in part by the US National Human Genome Research Institute/National Heart Blood Lung Institute jointly funded Baylor-Hopkins Center for Mendelian Genomics (UM1HG006542), by the National Institutes of Health (NIH) (5R01 HD039056, 5R01 HL091771), by the Genomic Research Elucidates the Genetics of Rare Disease (GREGoR) Program (U01 HG011758) to J.E.P., J.R.L., and R.A.G., and by the National Institute of Neurological Disorders and Stroke (NINDS R35 NS105078) to J.R.L. J.X. was supported by Simons Foundation pilot award (AGT011737). J.S. was supported by the Genomic Research Elucidates the Genetics of Rare Disease (GREGoR) Program (U01 HG011758). S.J was supported by the University of Colorado School of Medicine Translational Research Scholars Program, Simons Foundation pilot award (AGT011737), and the National Institutes of Health grant R35GM133433. Z.C.-A. was supported by the TOPMed NHLBI Fellowship, Simons Foundation pilot award (AGT011737) and the Genomic Research Elucidates the Genetics of Rare Disease (GREGoR) Program (U01 HG011758). S.S.Y. is a Partner Faculty Member of the GREGoR Consortium. The content is solely the responsibility of the authors and does not necessarily represent the official views of the NIH. Full acknowledgments are provided in the supplemental information.

## Author Contributions

I.E. and J.S. contributed equally to this work. Conceptualization: Z.C.-A. and S.J.; Methodology: I.E., J.S., M.C., J.X., and Z.C.-A.; Formal analysis: I.E., J.S., M.C., and J.X.; Resources: P.O., M.D., T.B.-Y., J.K., L.M., M.T., S.S.Y., D.C., J.E.P., R.A.G., E.B., A.P.R., P.S.d.V., A.C.M., C.A.S., J.R.L., C.M.B.C., S.M., S.J., and Z.C.-A.; Writing – original draft: I.E., J.S., S.J., and Z.C.-A.; Writing – review & editing: all authors; Supervision: Z.C.-A. and S.J.; Funding acquisition: Z.C.-A., S.J., E.B., R.A.G., J.R.L., and A.C.M. All co-authors read and approved the final manuscript. We are aware of other work developed by Dr. Steven E Brenner and we would like to thank him for the opportunity to co-submit our manuscripts.

## Declaration of generative AI and AI-assisted technologies

During the preparation of this work, the authors used Claude.ai to assist with language editing and to enhance readability. Following its use, the authors carefully reviewed and revised the content as necessary and assume full responsibility for the content of the publication.

## Supplemental information

Figures S1–S13, Tables S1 and S2. Supplemental Acknowledgments.

## Web resources

TOPMed: https://topmed.nhlbi.nih.gov/topmed-whole-genome-sequencing-methods-freeze-9 gnomAD: https://gnomad.broadinstitute.org/;

ClinVar: https://www.ncbi.nlm.nih.gov/clinvar/;

GTEx: https://www.gtexportal.org/;

GREGoR: https://gregorconsortium.org/data;

EJC RIP-Seq data: https://www.ncbi.nlm.nih.gov/geo/query/acc.cgi?acc=GSE41154;

ATtRACT database: https://attract.cnic.es/;

MEME Suite: https://meme-suite.org/

SnpEff https://pcingola.github.io/SnpEff/snpeff/inputoutput/

GENCODE: https://www.gencodegenes.org/human/release_26.html

GATK ASEReadCounter (v3): https://gatk.broadinstitute.org/hc/en-us/articles/360037428291-ASEReadCounter

Ensembl BioMart: https://www.ensembl.org/biomart

