## Supplemental Figures 1-13 for "Unveiling the Hidden Rules: Enhancing NMD Prediction for Protein-Truncating Variants"

### Supplemental Information

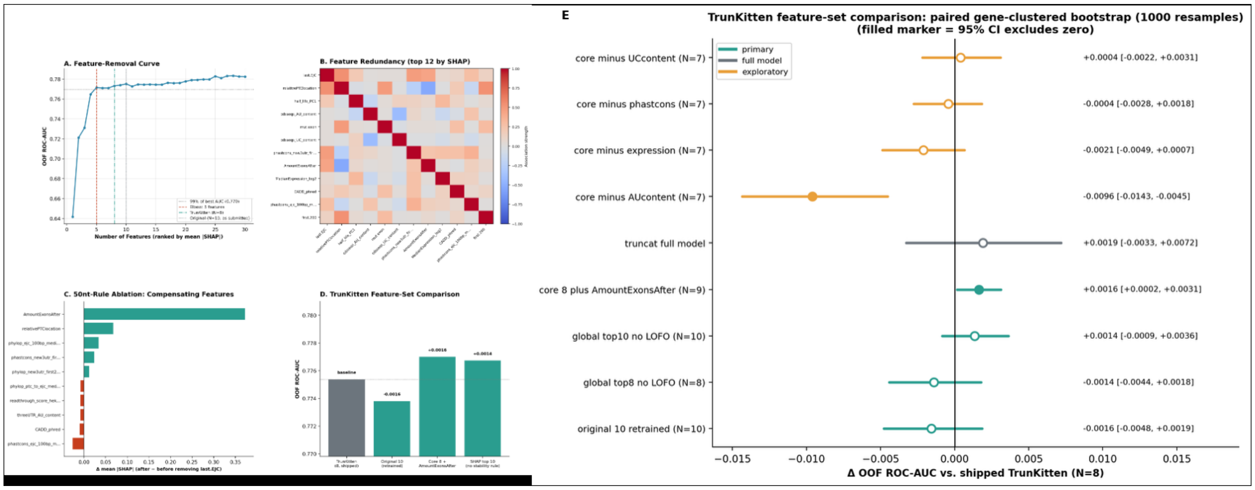


**Figure S1. Feature-ablation, redundancy, and compensation analyses for TrunCat and TrunKitten.** All models were retrained under the same 5-fold gene-grouped cross-validation used for TrunCat. (A) Out-of-fold (OOF) ROC-AUC as a function of the number of features included (1–30), ranked by mean |SHAP|; vertical lines mark the five-feature elbow, the eight-feature TrunKitten, and the original ten-feature set, and the horizontal dotted line marks 99% of the best AUC. (B) Pairwise association among the 12 highest-ranked features by mean |SHAP|. (C) Change in mean |SHAP| for each feature after removing last.EJC (the 50-nt-rule feature) and retraining; AmountExonsAfter absorbs most of the signal. (D) OOF ROC-AUC of the eight-feature TrunKitten compared with the original ten features retrained, the eight TrunKitten features plus AmountExonsAfter, and the top ten features by mean |SHAP| selected without the stability rule. (E) Difference in OOF ROC-AUC relative to TrunKitten for alternative feature sets (primary), the full TrunCat model, and TrunKitten with single features removed (exploratory), estimated by gene-clustered paired bootstrap (1,000 resamples); horizontal bars show 95% confidence intervals, and filled markers indicate intervals that exclude zero.

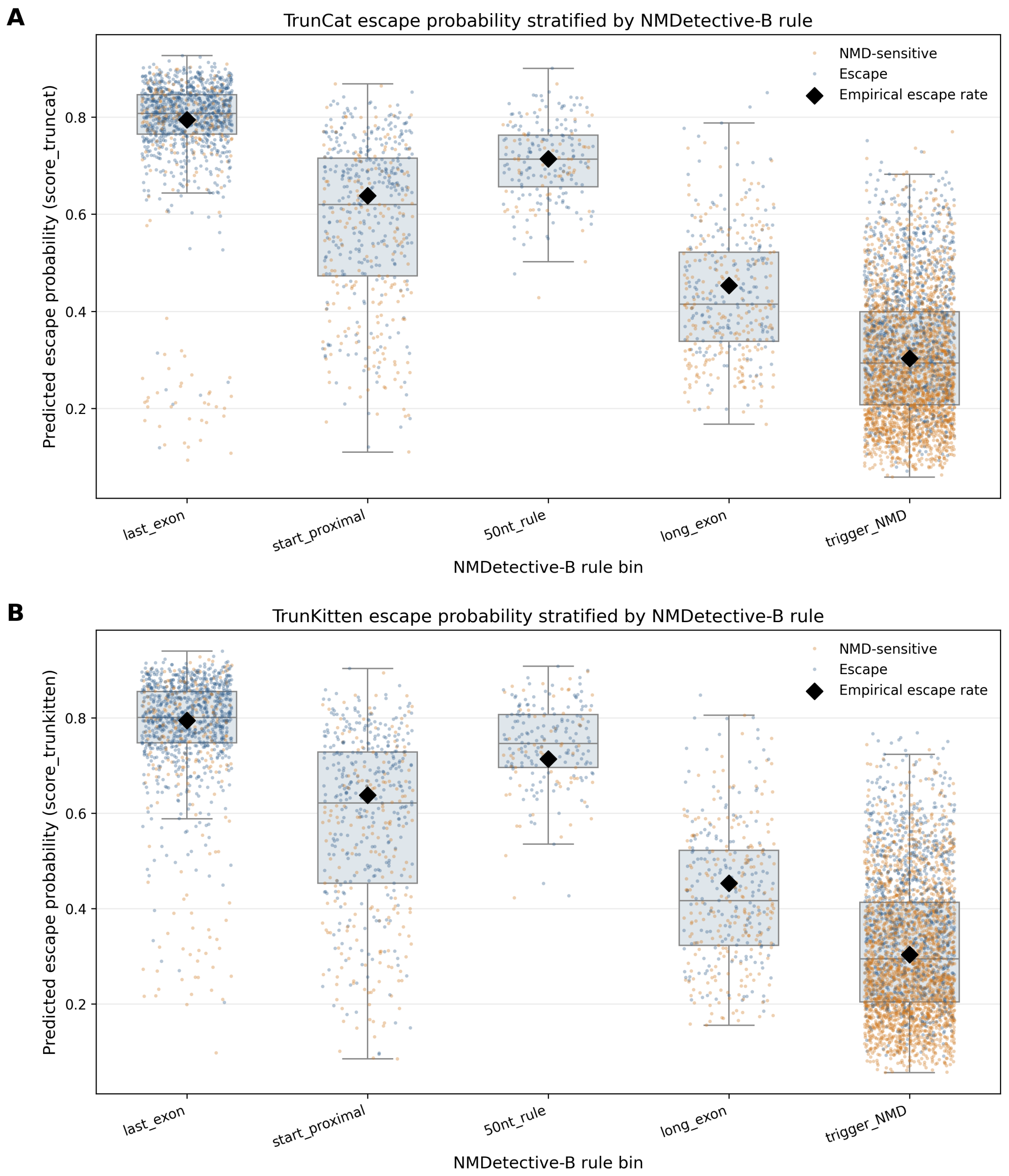


**Figure S2. TrunCat and TrunKitten predictions stratified by NMDetective-B rule bins.** (A) Distribution of out-of-fold NMD-escape probabilities predicted by the retrained TrunCat model (score_truncat) for the 5,749 TOPMed variants, stratified by NMDetective-B rule bin (last_exon, n = 1,123; start_proximal, n = 517; 50nt_rule, n = 217; long_exon, n = 399; trigger_NMD, n = 3,493). Boxes show the median and interquartile range; points are individual variants colored by observed class (NMD-sensitive, orange; escape, blue); black diamonds mark the empirical escape rate within each bin. (B) Same as (A) for the 8-feature TrunKitten model (score_trunkitten). In both models, the median predicted escape probability closely tracks the empirical escape rate in every rule bin, from highest in the last_exon bin to lowest in the trigger_NMD bin, while the wide spread of predictions within bins reflects variant-level differences that the rule assignment alone does not capture.

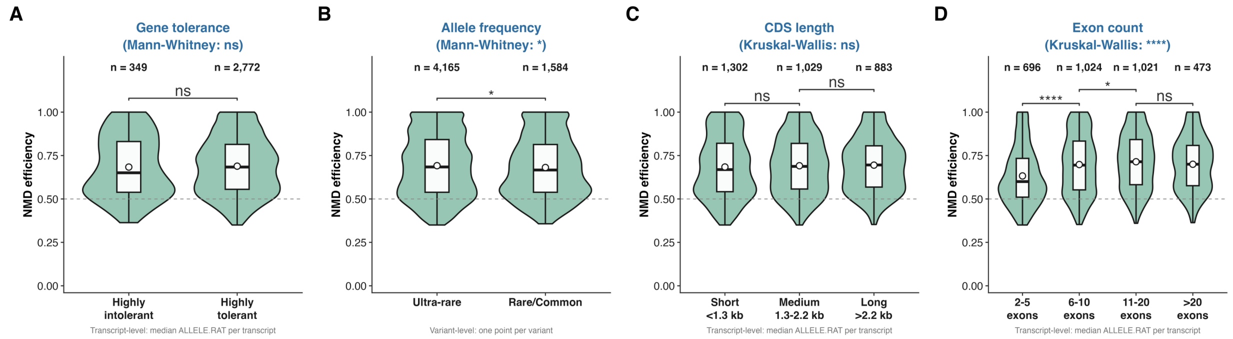


**Figure S3. NMD efficiency stratified by gene tolerance, allele frequency, CDS length, and exon count.** Distributions of NMD efficiency for TOPMed stop-gain variants with allele ratio ≥ 0.35, across (A) gene tolerance (highly intolerant, pLI ≥ 0.65, n = 349 transcripts; highly tolerant, pLI < 0.35, n = 2,772 transcripts), (B) allele frequency (ultra-rare, n = 4,165 variants; rare/common, n = 1,584 variants), (C) CDS length (short, <1.3 kb, n = 1,302; medium, 1.3–2.2 kb, n = 1,029; long, >2.2 kb, n = 883 transcripts), and (D) exon count (2–5 exons, n = 696; 6–10, n = 1,024; 11–20, n = 1,021; >20, n = 473 transcripts). Panels A, C, and D are transcript-level (median NMD efficiency per transcript); panel B is variant-level, because allele frequency is a property of the variant rather than the transcript. Violins show the distribution, boxes the median and interquartile range, and white circles the mean; the dashed line marks NMD efficiency = 0.5. The overall test for each panel is given in its title: Mann-Whitney U test for two-group comparisons (A, B) and Kruskal-Wallis test for multi-group comparisons (C, D). Brackets show pairwise Wilcoxon rank-sum tests between adjacent groups with Benjamini-Hochberg correction (ns, not significant; *P < 0.05; ****P < 0.0001). Gene tolerance and CDS length show no significant association with NMD efficiency, and allele frequency shows a small but significant difference (P < 0.05). Exon count shows a strong association (Kruskal-Wallis P < 0.0001), driven mainly by lower NMD efficiency in transcripts with 2–5 exons.

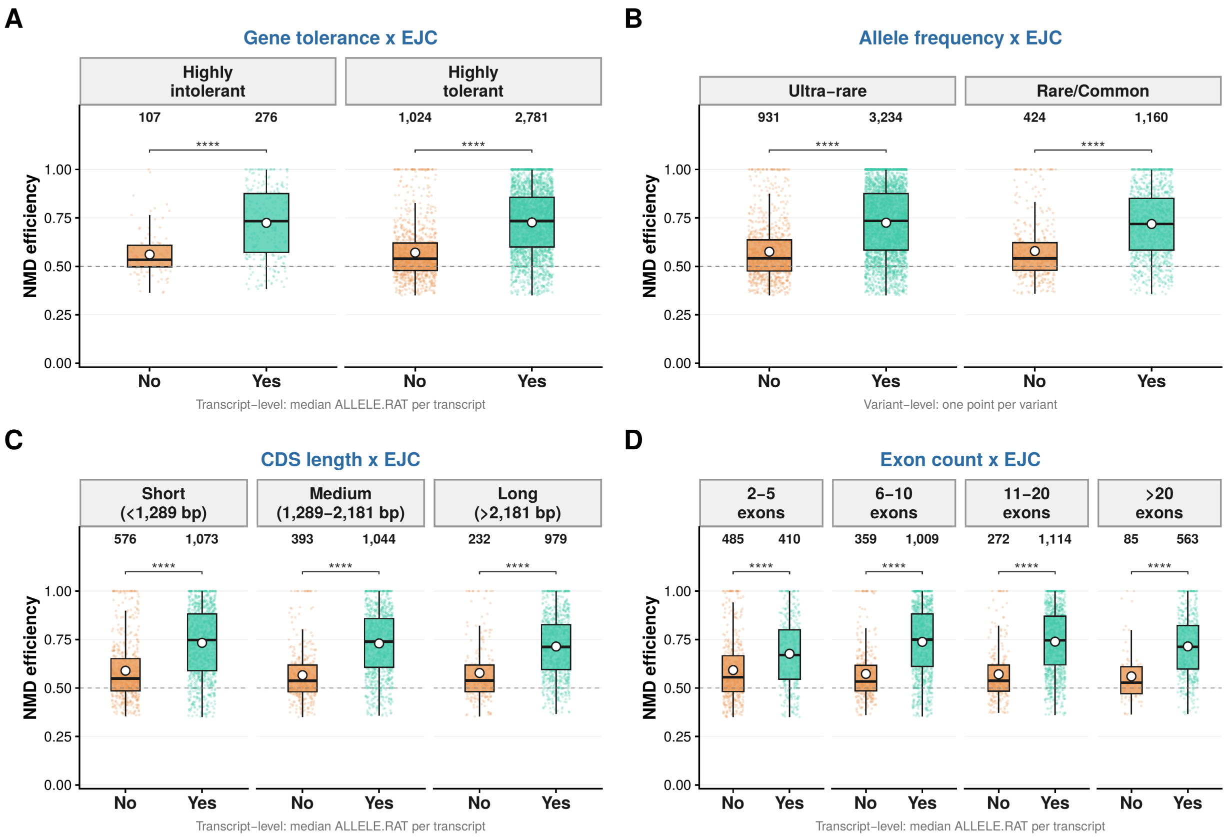


**Figure S4. The canonical downstream-EJC effect is robust across gene constraint, allele frequency, and transcript architecture.** NMD efficiency of PTCs with (Yes) and without (No) a downstream exon junction complex (EJC), stratified by (A) gene tolerance (highly intolerant, pLI $\geq$ 0.65, n = 107 No / 276 Yes; highly tolerant, pLI < 0.35, n = 1,024 No / 2,781 Yes), (B) allele frequency (ultra-rare, n = 931 No / 3,234 Yes; rare/common, n = 424 No / 1,160 Yes), (C) CDS length (short, <1,289 bp, n = 576 No / 1,073 Yes; medium, 1,289–2,181 bp, n = 393 No / 1,044 Yes; long, >2,181 bp, n = 232 No / 979 Yes), and (D) exon count (2–5 exons, n = 485 No / 410 Yes; 6–10, n = 359 No / 1,009 Yes; 11–20, n = 272 No / 1,114 Yes; >20, n = 85 No / 563 Yes). Variants with allele ratio $\geq$ 0.35 were included. Panel B shows one point per variant; panels A, C, and D show the median NMD efficiency per transcript, computed separately for each EJC status and allele-frequency class. Boxes show the median and interquartile range and white circles the mean; the dashed line marks NMD efficiency = 0.5. In every stratum, NMD efficiency is significantly higher when a downstream EJC is present (Wilcoxon rank-sum test, ****$P < 0.0001$).

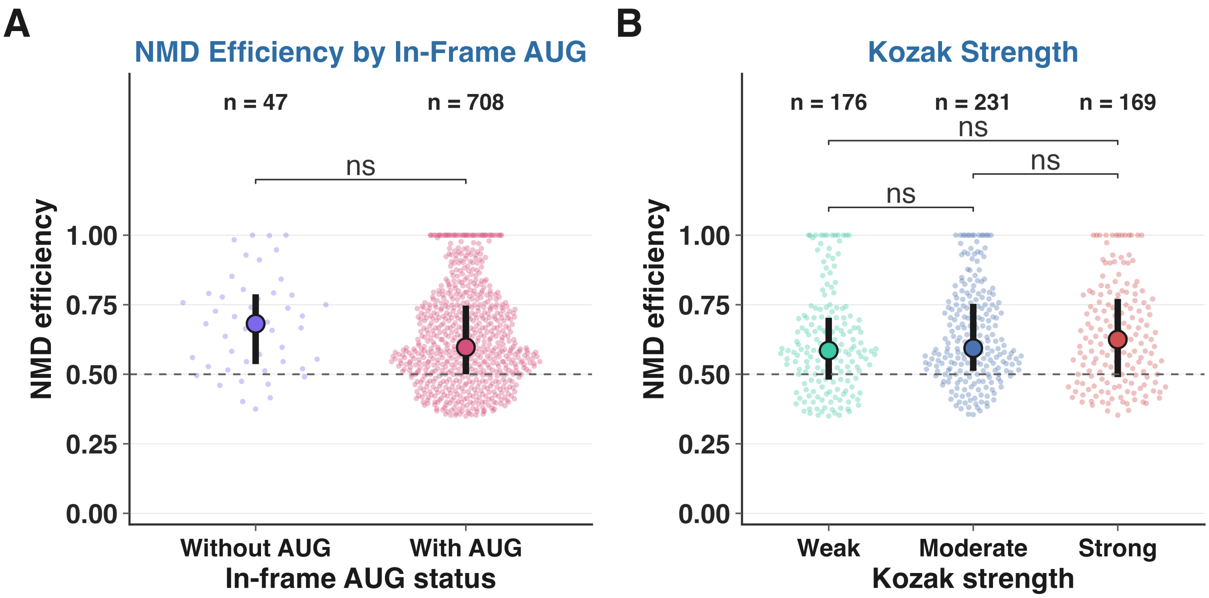


**Figure S5. In-frame downstream AUG and Kozak context of start-proximal PTCs.** Variant-level NMD efficiency for nonsense variants with allele ratio ≥ 0.35 whose PTC lies in the first 200 bp of the coding sequence. (A) NMD efficiency of variants without (n = 47) and with (n = 708) at least one in-frame downstream AUG; the difference is not significant (median 0.68 vs. 0.60; P = 0.08). (B) NMD efficiency by Kozak-context strength of the in-frame downstream AUG (weak, n = 176; moderate, n = 231; strong, n = 169); all pairwise comparisons are not significant. Points are individual variants, large dots mark the median and bars the interquartile range; the dashed line marks NMD efficiency = 0.5. Two-sided Wilcoxon rank-sum tests (ns, not significant).

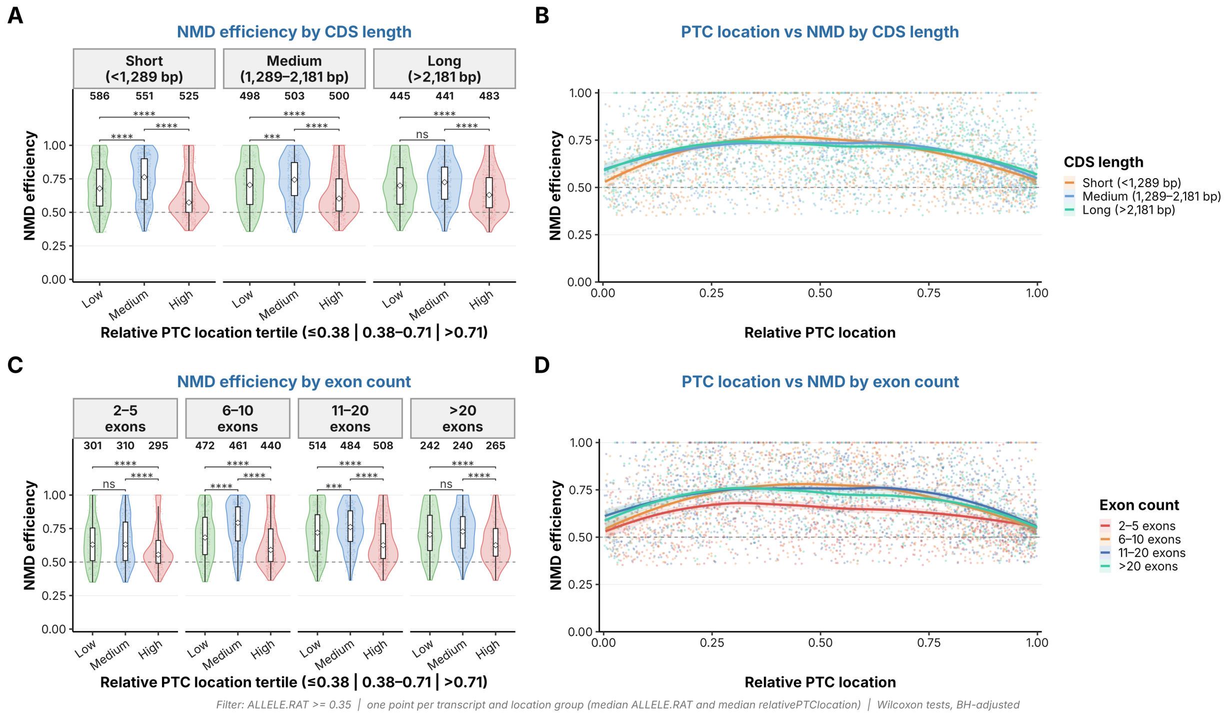


**Figure S6. Relative PTC location and NMD efficiency stratified by transcript architecture.** (A) NMD efficiency by relative PTC location tertile, stratified by CDS length (short, <1,289 bp; medium, 1,289–2,181 bp; long, >2,181 bp). Location groups are tertiles of relative PTC location across all variants: low (≤0.38), medium (0.38–0.71), and high (>0.71), each containing one third of the variants. Each point is the median NMD efficiency of the variants of one transcript within one location group; n is shown above each violin (short, 586 / 551 / 525; medium, 498 / 503 / 500; long, 445 / 441 / 483). (B) Relative PTC location versus NMD efficiency with LOESS smoothers (shaded bands, 95% confidence intervals) for each CDS-length group. (C) As in (A), stratified by exon count (2–5 exons, n = 301 / 310 / 295; 6–10, n = 472 / 461 / 440; 11–20, n = 514 / 484 / 508; >20, n = 242 / 240 / 265). (D) As in (B), stratified by exon count. NMD efficiency is highest for PTCs at intermediate relative positions in every CDS-length and exon-count group: PTCs in the high-location tertile have lower NMD efficiency than those in the medium tertile in all strata ($P < 0.0001$), and lower than those in the low tertile in all strata ($P < 0.0001$). PTCs in the low tertile are less efficient than those in the medium tertile in short and medium-length CDSs and in transcripts with 6–20 exons, but not in long CDSs or in transcripts with 2–5 or >20 exons. The pattern is weakest in transcripts with 2–5 exons, whose LOESS curve is the flattest. Variants with allele ratio ≥ 0.35 were included; pairwise Wilcoxon rank-sum tests with Benjamini-Hochberg correction (ns, not significant; ***$P < 0.001$; ****$P < 0.0001$).

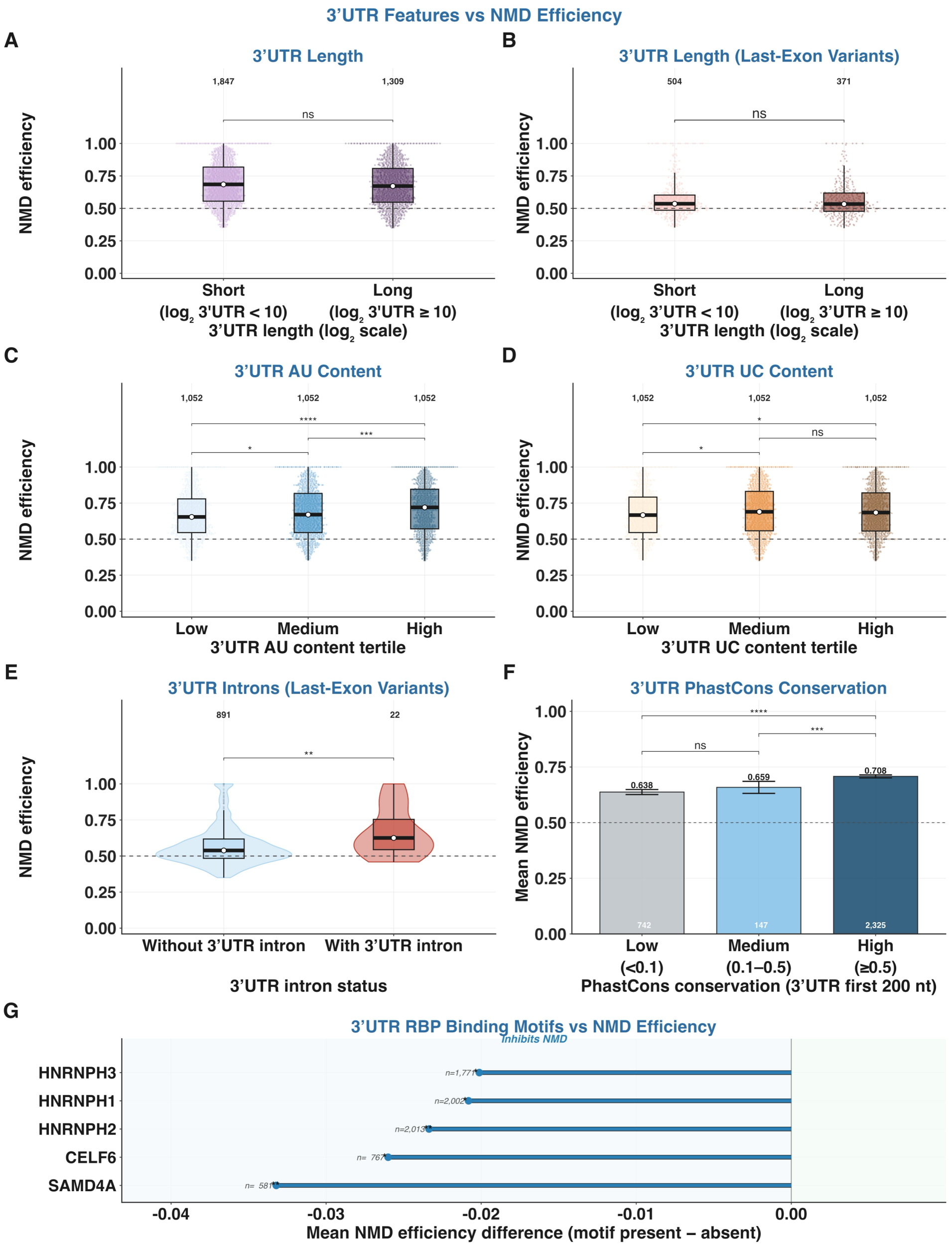


**Figure S7. 3′UTR features and their relationship to NMD efficiency.** Transcript-level NMD efficiency (median per transcript) for variants with allele ratio ≥ 0.35. (A) 3′UTR length (short, $\log_2 < 10$, n = 1,847; long, $\log_2 \geq 10$, n = 1,309) for all variants; ns. (B) Same comparison restricted to last-exon variants (short, n = 504; long, n = 371); ns. (C) 3′UTR AU-content tertiles (n = 1,052 each); NMD efficiency increases with AU content (low vs medium, $P < 0.05$; medium vs high, $P < 0.001$; low vs high, $P < 0.0001$). (D) 3′UTR UC-content tertiles (n = 1,052 each); low vs medium, $P < 0.05$; low vs high, $P < 0.05$; medium vs high, ns. (E) Presence or absence of a 3′UTR intron among last-exon variants (without, n = 891; with, n = 22); $P < 0.01$, based on a small group of intron-containing transcripts. (F) Mean NMD efficiency (± 95% confidence interval) by PhastCons conservation of the first 200 nt of the 3′UTR (low, <0.1, n = 742, mean 0.638; medium, 0.1–0.5, n = 147, mean 0.659; high, ≥0.5, n = 2,325, mean 0.708); low vs high, $P < 0.0001$;

medium vs high, $P < 0.001$; low vs medium, ns. (G) Bonferroni-significant RBP binding motifs in the 3′UTR, ranked by the difference in mean NMD efficiency between transcripts with and without the motif. All five recovered motifs (SAMD4A, CELF6, HNRNPH2, HNRNPH1, and HNRNPH3) are associated with lower NMD efficiency. Comparisons in A–F use Wilcoxon rank-sum tests (ns, not significant; *$P < 0.05$; **$P < 0.01$; ***$P < 0.001$; ****$P < 0.0001$).

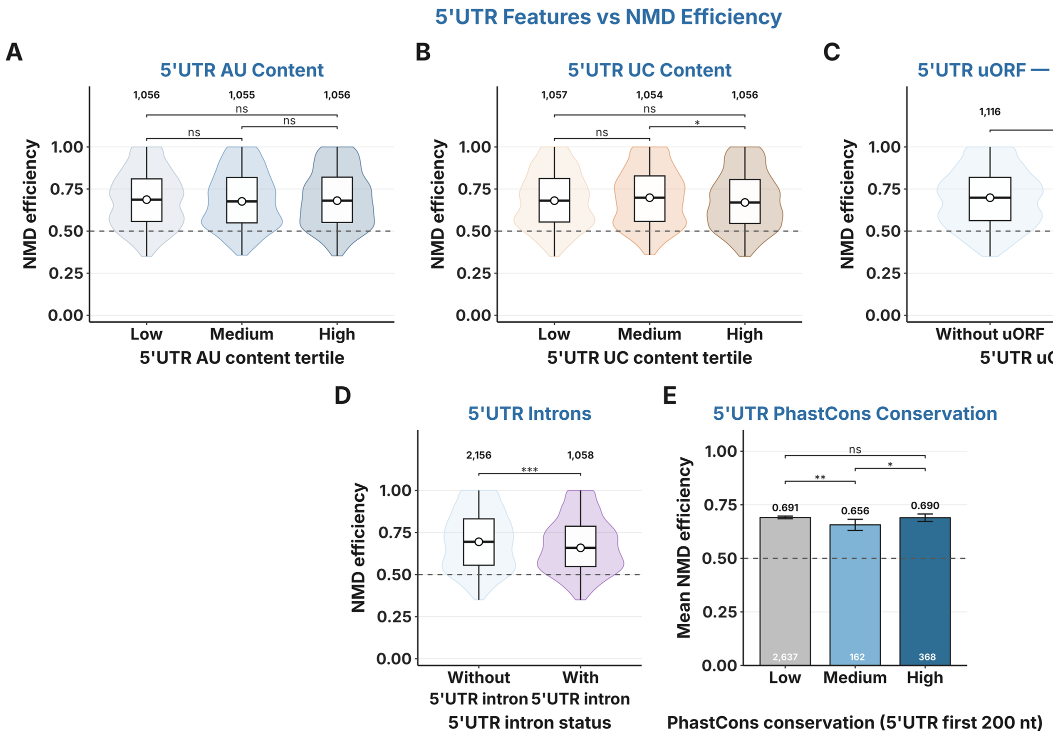


**Figure S8. 5′UTR features show weak or no association with NMD efficiency.** Transcript-level NMD efficiency (median per transcript) for variants with allele ratio ≥ 0.35. (A) 5′UTR AU-content tertiles (n = 1,056 / 1,055 / 1,056); all comparisons ns. (B) 5′UTR UC-content tertiles (n = 1,057 / 1,054 / 1,056); medium vs high, $P < 0.05$; other comparisons ns. (C) Presence or absence of an upstream open reading frame (uORF) in the 5′UTR (without, n = 1,116; with, n = 2,051 transcripts; transcripts without an annotated 5′UTR excluded); ns. (D) Presence or absence of a 5′UTR intron (without, n = 2,156; with, n = 1,058); $P < 0.001$, with slightly lower NMD efficiency in transcripts that contain a 5′UTR intron. (E) Mean NMD efficiency (± 95% confidence interval) by PhastCons conservation of the first 200 nt of the 5′UTR (low, <0.1, n = 2,637, mean 0.691; medium, 0.1–0.5, n = 162, mean 0.656; high, ≥0.5, n = 368, mean 0.690); low vs medium, $P < 0.01$; medium vs high, $P < 0.05$; low vs high, ns. Wilcoxon rank-sum tests (ns, not significant; *$P < 0.05$; **$P < 0.01$; ***$P < 0.001$). Apart from these modest differences, 5′UTR architecture is not a major determinant of NMD efficiency in germline PTC variants.

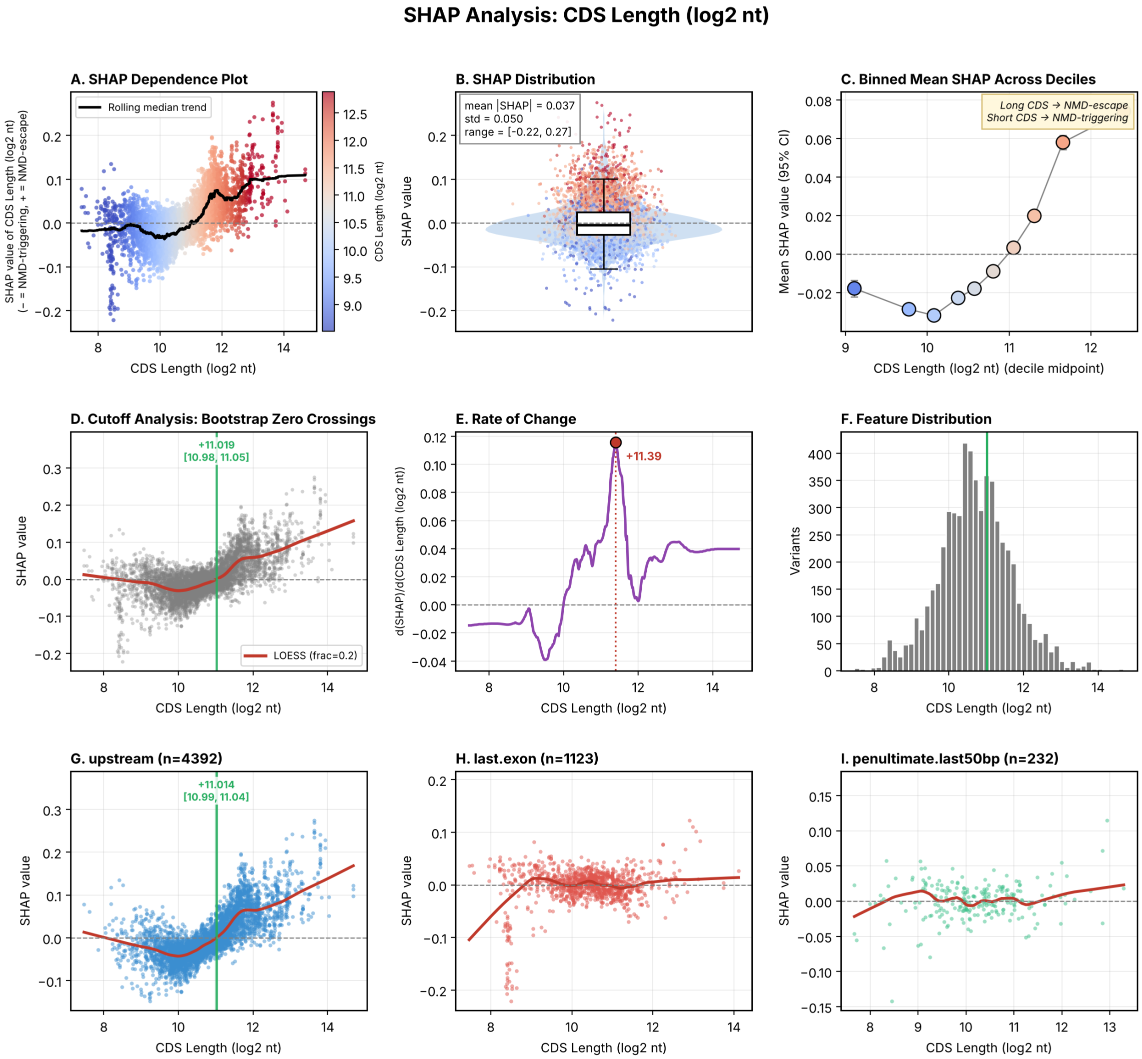


**Figure S9. SHAP analysis of CDS length.** SHAP values are cross-validation-averaged out-of-fold values from the retrained TrunCat model (n = 5,747 variants). (A) SHAP dependence plot of per-variant SHAP values for CDS length ($\log_2$ nt), coloured by CDS length, with a rolling-median trend (black). Negative SHAP values push the prediction toward NMD-triggering and positive values toward NMD-escape. Mean |SHAP| = 0.037, SD 0.050, range [−0.22, 0.27]. (B) Distribution of SHAP values for CDS length. (C) Mean SHAP value (95% CI) across CDS-length deciles, shifting from negative (NMD-triggering) at short CDS to positive (NMD-escape) at long CDS. (D) Bootstrap zero-crossing analysis of the LOESS fit (frac = 0.2; 500 bootstrap resamples) identifies the transition at $\log_2$ CDS = 11.02 (95% CI 10.98–11.05). (E) The rate of change d(SHAP)/d($\log_2$ CDS) peaks at $\log_2$ CDS = 11.39. (F) Distribution of CDS lengths; the green line marks the zero crossing. (G–I) SHAP dependence stratified by EJC class: upstream (n = 4,392; crossing at 11.01 [10.99, 11.04]), last.exon (n = 1,123), and penultimate.last50bp (n = 232). In the last two classes the LOESS curve stays close to zero and no robust crossing is detected, indicating that CDS length contributes mainly for PTCs upstream of the last exon junction. Zero crossings are reported only where the LOESS curve departs from zero by at least 10% of the pooled curve's maximum on both sides and the crossing is recovered in at least 80% of bootstrap resamples.

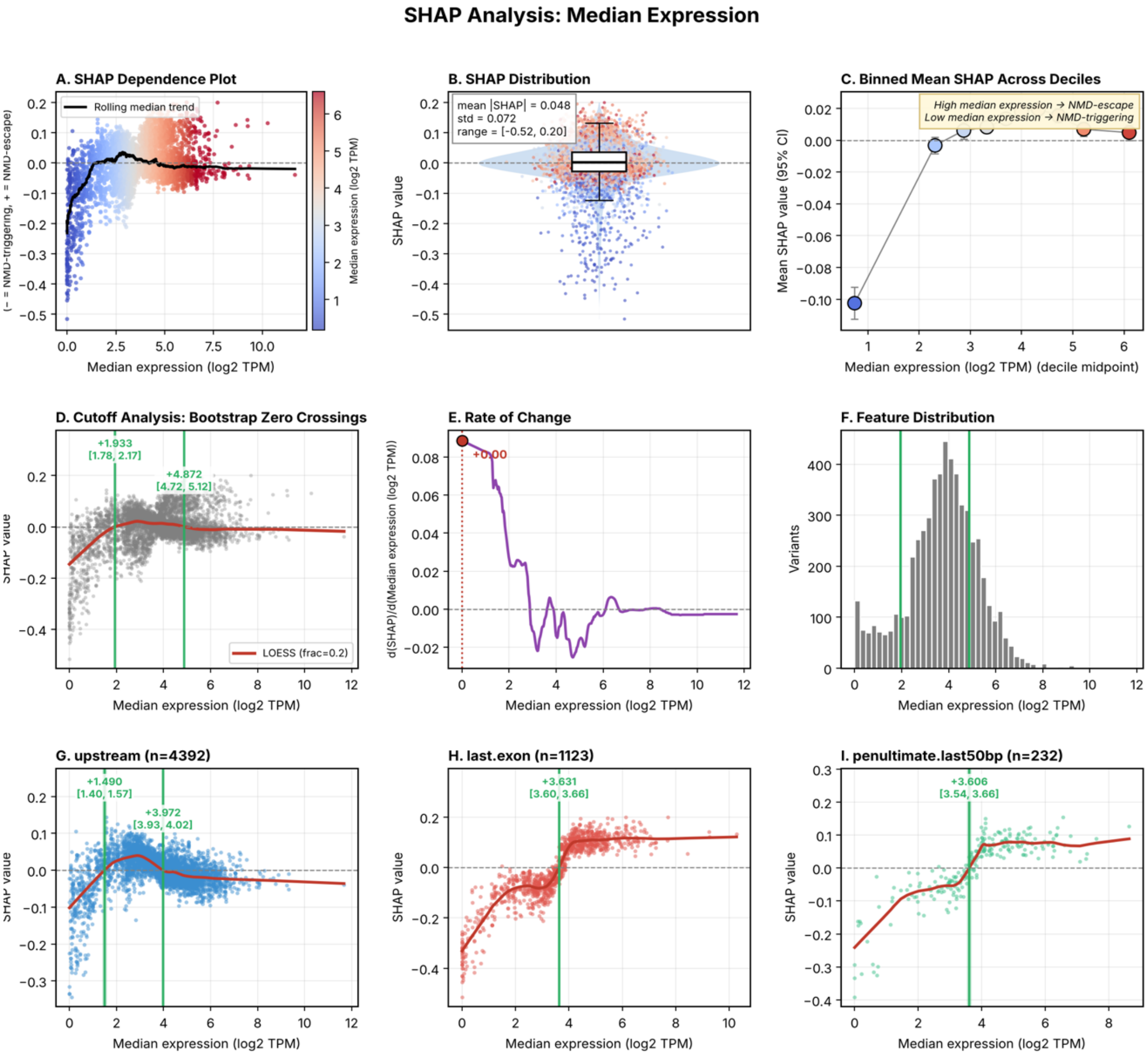


**Figure S10. SHAP analysis of median transcript expression.** SHAP values are cross-validation-averaged out-of-fold values from the retrained TruncCat model (n = 5,747 variants). (A) SHAP dependence plot of per-variant SHAP values for median expression ($\log_2$ TPM), coloured by expression, with a rolling-median trend (black). Negative SHAP values push the prediction toward NMD-triggering and positive values toward NMD-escape. Mean |SHAP| = 0.048, SD 0.072, range [−0.52, 0.20]. (B) Distribution of SHAP values for median expression. (C) Mean SHAP value (95% CI) across expression deciles: strongly negative (NMD-triggering) in the lowest decile and slightly positive (NMD-escape) at higher expression. (D) Bootstrap zero-crossing analysis of the LOESS fit (frac = 0.2; 500 bootstrap resamples) identifies pooled crossings at $\log_2$ TPM = 1.93 (95% CI 1.78–2.17) and 4.87 (95% CI 4.72–5.12). (E) The rate of change d(SHAP)/d(median expression) is largest at the lowest expression values ($\log_2$ TPM ≈ 0). (F) Distribution of median-expression values; green lines mark the zero crossings. (G–I) SHAP dependence stratified by EJC class: upstream (n = 4,392; crossings at 1.49 [1.40, 1.57] and 3.97 [3.93, 4.02]), last.exon (n = 1,123; crossing at 3.63 [3.60, 3.66]), and penultimate.last50bp (n = 232; crossing at 3.61 [3.54, 3.66]). In last-exon and penultimate-last-50-bp variants, expression above $\log_2$ TPM $\approx$ 3.6 consistently pushes predictions toward NMD escape. Zero crossings are reported only where the LOESS curve departs from zero by at least 10% of the pooled curve's maximum on both sides and the crossing is recovered in at least 80% of bootstrap resamples.

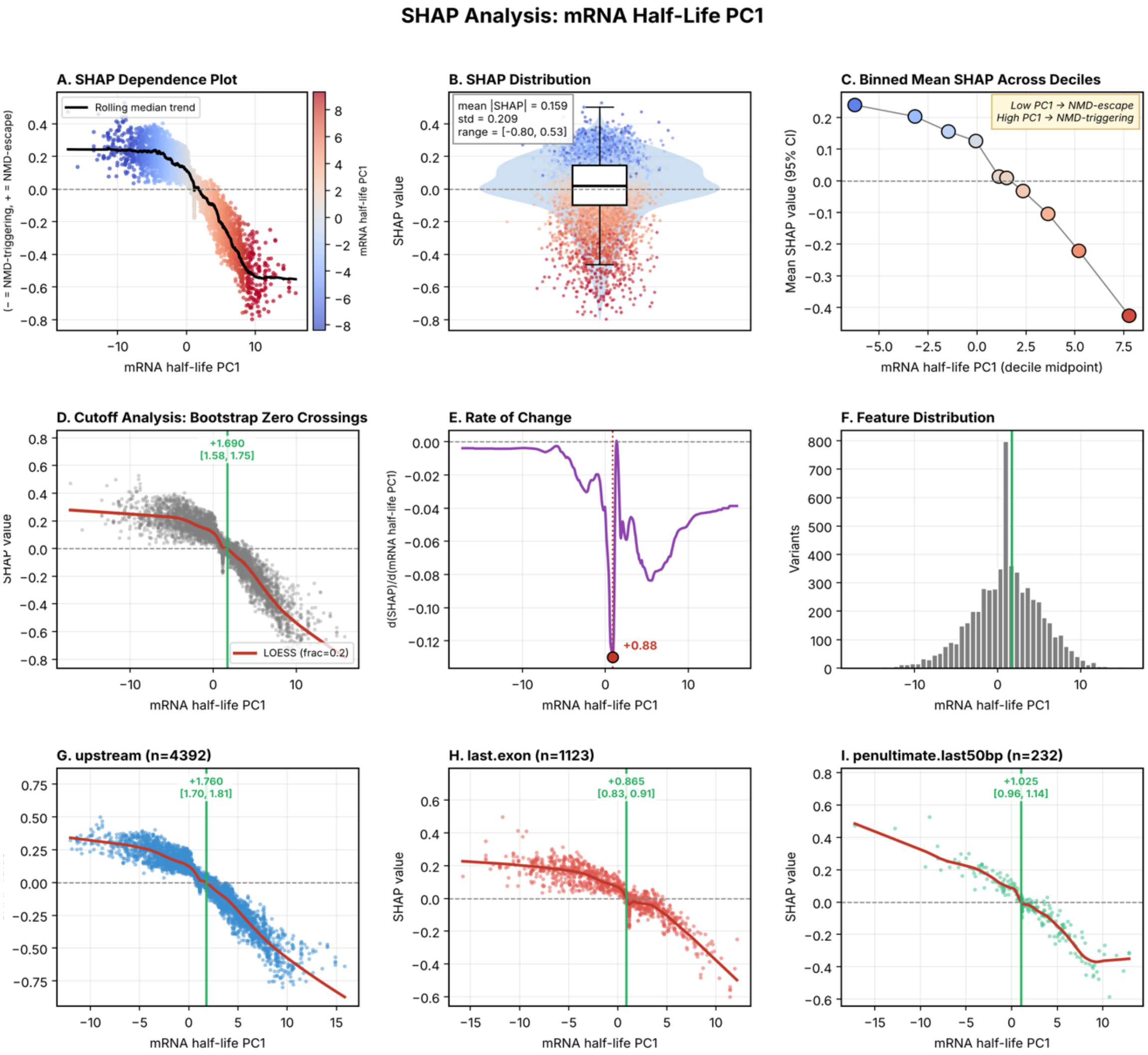


**Figure S11. SHAP analysis of mRNA half-life PC1.** SHAP values are cross-validation-averaged out-of-fold values from the retrained TrunCat model (n = 5,747 variants). (A) SHAP dependence plot of per-variant SHAP values for mRNA half-life PC1, coloured by PC1, with a rolling-median trend (black). Negative SHAP values push the prediction toward NMD-triggering and positive values toward NMD-escape. Mean |SHAP| = 0.159, SD 0.209, range [−0.80, 0.53]. (B) Distribution of SHAP values for mRNA half-life PC1. (C) Mean SHAP value (95% CI) across PC1 deciles, shifting monotonically from positive (NMD-escape) at low PC1 to negative (NMD-triggering) at high PC1. (D) Bootstrap zero-crossing analysis of the LOESS fit (frac = 0.2; 500 bootstrap resamples) identifies the pooled crossing at PC1 = 1.69 (95% CI 1.58–1.75). (E) The rate of change d(SHAP)/d(PC1) reaches its largest magnitude at PC1 = 0.88. (F) Distribution of PC1 values; the green line marks the zero crossing. (G–I) SHAP dependence stratified by EJC class: upstream (n = 4,392; crossing at 1.76 [1.70, 1.81]), last.exon (n = 1,123; crossing at 0.87 [0.83, 0.91]), and penultimate.last50bp (n = 232; crossing at 1.03 [0.96, 1.14]). The monotonic SHAP profile is preserved in all three EJC classes, indicating that mRNA half-life PC1 is a robust, continuous determinant of NMD efficiency at the model level, consistent with the coupling between NMD and translation. Zero crossings are reported only where the LOESS curve departs from zero by at least 10% of the pooled curve's maximum on both sides and the crossing is recovered in at least 80% of bootstrap resamples.

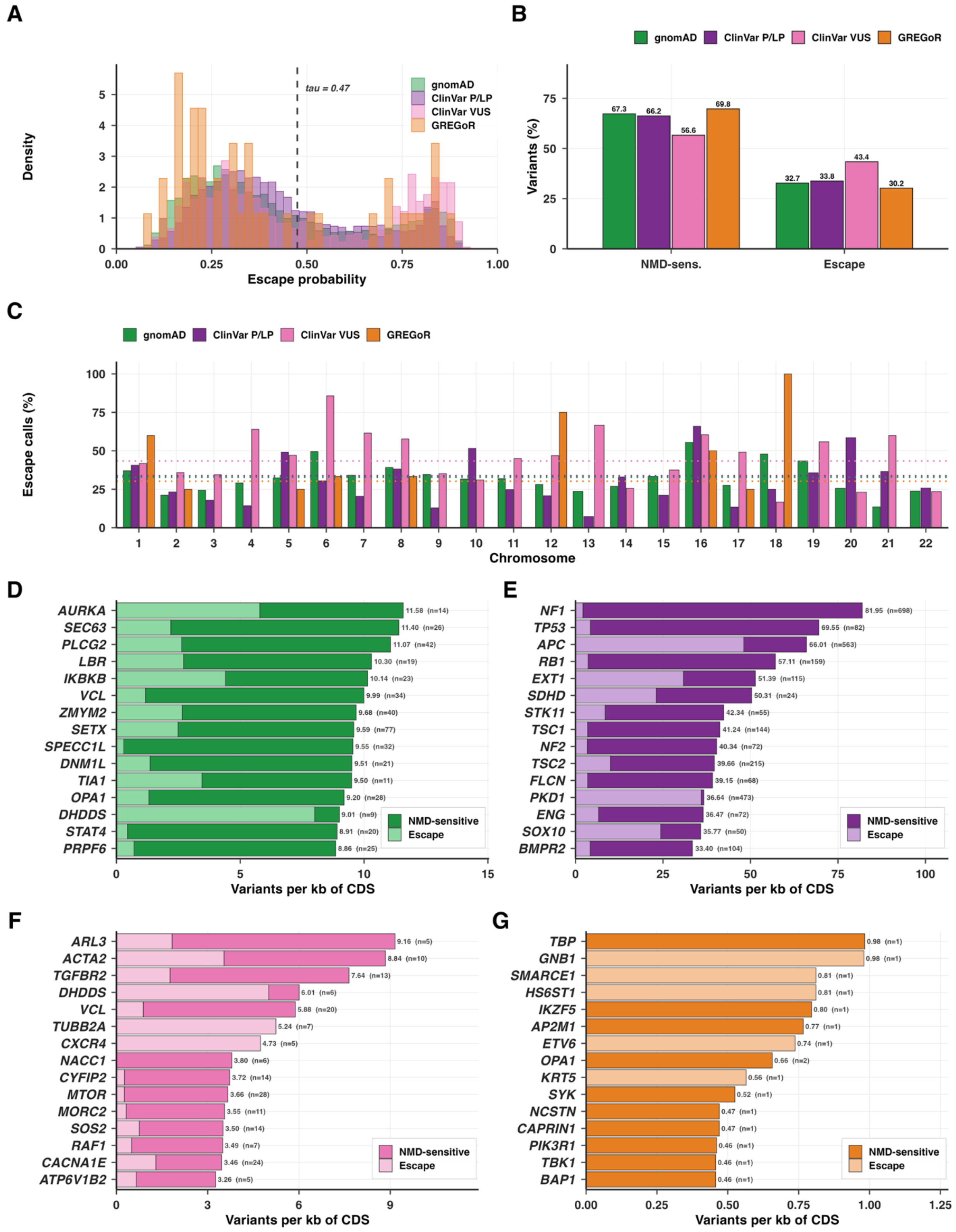


**Figure S12. Population- and clinical-scale escape-probability landscape across gnomAD, ClinVar P/LP, ClinVar VUS, and the GREGoR rare-disease cohort, restricted to autosomal-**

**dominant, highly LoF-intolerant genes (LOEUF < 0.6, pLI ≥ 0.9; 778 genes).** Escape probabilities are from the retrained TrunCat model. (A) Normalized escape-probability distributions for gnomAD, ClinVar pathogenic/likely pathogenic (P/LP), ClinVar variants of uncertain significance (VUS), and the GREGoR rare-disease cohort within the autosomal-dominant intolerant-gene subset; the dashed line marks the classification threshold $\tau = 0.47$. (B) Class composition (% NMD-sensitive vs. % escape) in each dataset within this subset: gnomAD 67.3/32.7, ClinVar P/LP 66.2/33.8, ClinVar VUS 56.6/43.4, GREGoR 69.8/30.2. (C) Per-chromosome predicted escape fractions; dotted lines indicate dataset means. (D–G) Top 15 genes by variant density, the number of called variants per kilobase of coding sequence (CDS), in gnomAD (D), ClinVar P/LP (E), ClinVar VUS (F), and GREGoR (G). CDS lengths are from the MANE Select transcript in the gnomAD v4.1 constraint table. Genes with at least five variants were ranked in D–F; because few GREGoR variants fall in this gene subset, all GREGoR genes were ranked in G. Each bar is split into predicted NMD-sensitive (dark shading) and NMD-escape (light shading) calls, and the total per-gene density and variant count (n) are shown to the right of each bar. Normalizing by CDS length removes the gene-size bias that dominates raw variant counts and surfaces loci that are mutationally dense per coding base rather than simply long. Within this constrained-gene subset, gnomAD is led by *AURKA*, *SEC63*, *PLCG2*, *LBR*, and *IKBKB*; ClinVar P/LP by *NF1*, *TP53*, *APC*, *RB1*, and *EXT1*; ClinVar VUS by *ARL3*, *ACTA2*, *TGFBR2*, *DHDDS*, and *VCL*; and the GREGoR cohort by *TBP*, *GNB1*, *SMARCE1*, *HS6ST1*, and *IKZF5*.

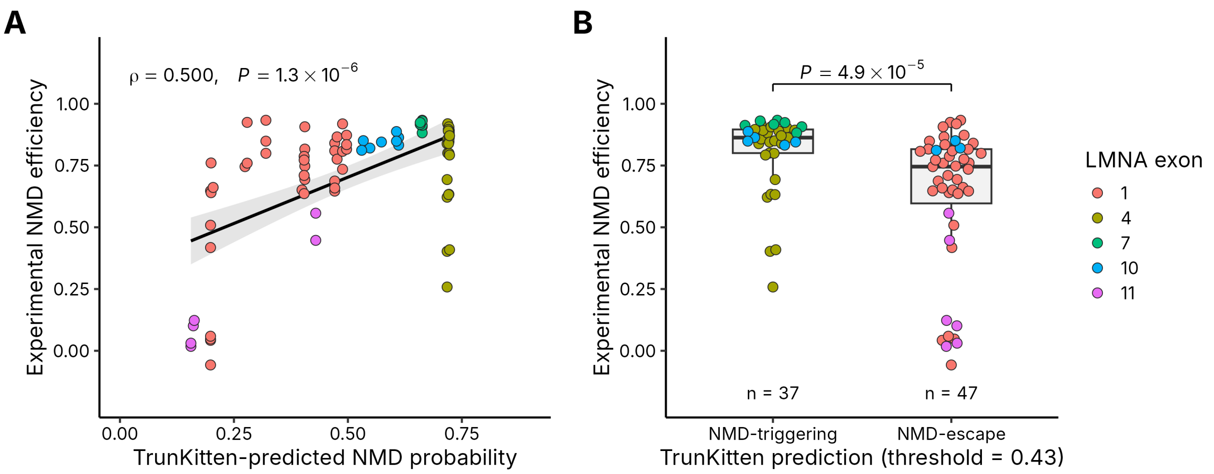


**Figure S13. TrunKitten predictions are concordant with experimentally measured NMD efficiency in LMNA.** Experimental NMD efficiency from the LMNA saturation genome editing (SGE) dataset of Cortázar et al. (2025) for 84 stop-gain variants that can arise from the endogenous codon by a single-nucleotide substitution and passed the experimental variability filter (SD ≤ 0.2), compared with predictions from the eight-feature TrunKitten model applied without retraining. (A) TrunKitten-predicted NMD probability (1 − escape probability) versus experimental NMD efficiency; line, linear fit with 95% confidence band; Spearman ρ = 0.500, P = 1.3 × $10^{-6}$. (B) Experimental NMD efficiency of variants predicted to trigger NMD (n = 37) or to escape NMD (n = 47) at the TrunKitten decision threshold (escape probability 0.43); median 0.86 vs. 0.75; two-sided Wilcoxon rank-sum test, P = 4.9 × $10^{-5}$. Boxes show the median and interquartile range. Points are colored by LMNA exon; exon 11 is the penultimate exon.

**TOPMed ASE cohort**

| cohort | size | cohort | size |
|---|---|---|---|
| COPDGene | 709 | FHS_Levy | 201 |
| FHS_Ramachandran | 592 | GALAII | 1897 |
| LTRC | 1360 | MESA | 1985 |
| SAGE | 705 | SPIROMICS | 1578 |
| WHI | 1279 | | |

| Tissue types | Size | Tissue type | Size |
|---|---|---|---|

| | | | |
|---|---|---|---|
| Whole Blood | 6,602 | T-cell | 368 |
| PBMC | 1,265 | Monocyte | 352 |
| Lung | 1,360 | Nasal epithelial | 359 |

**Table S1**. **Composition of the TOPMed allele-specific expression (ASE) cohort.** (Top) Number of samples contributed by each constituent TOPMed study: COPDGene (709), FHS_Levy (201), FHS_Ramachandran (592), GALAII (1,897), LTRC (1,360), MESA (1,985), SAGE (705), SPIROMICS (1,578), and WHI (1,279). (Bottom) Distribution of the same samples across profiled tissue types: whole blood (6,602), PBMC (1,265), lung (1,360), T cell (368), monocyte (352), and nasal epithelial (359). Both breakdowns total 10,306 samples drawn from paired whole-genome sequencing and RNA-seq data, which constitute the ASE cohort used for NMD outcome calling.

| Filter | TOPMed | gnomAD | ClinVar | GREGoR |
|---|---|---|---|---|
| None | 11,145 | 285,776 | 57,394 | 6,681 |
| Single-exon removed | 10,583 | 267,924 | 56,155 | 6,019 |
| Low-expression genes removed | 6,892 | 137,859 | 30,130 | 2,550 |
| eGenes removed | 6,890 | 137,857 | 30,128 | 2,548 |
| Total reads count > 8 | 5,749 | -- | -- | -- |

**Table S2. Nonsense variant counts across successive quality-control filters in each dataset.** Number of nonsense (PTC-variants_ remaining after each sequential filtering step, shown separately for the TOPMed ASE cohort, gnomAD, ClinVar, and GREGoR. Starting from all called nonsense variants (None: TOPMed 11,145; gnomAD 285,776; ClinVar 57,394; GREGoR 6,681), variants were removed if they fell in single-exon transcripts (Single-exon removed: 10,583; 267,924; 56,155; 6,019), in low-expression genes (Low-expression genes removed: 6,892; 137,859; 30,130; 2,550), and in eGenes (eGenes removed: 6,890; 137,857; 30,128; 2,548). For TOPMed, a final filter retaining only variants with total read count > 8 yielded the 5,749 variants used for model training; this expression-based step was not applicable to gnomAD, ClinVar, or GREGoR (indicated by dashes), as these datasets lack the paired RNA-seq read depth required.

**Supplemental Acknowledgments**

The authors thank the TOPMed program and all study participants whose data made this work possible. This research was supported by funding from TOPMed program. Molecular data for TOPMed program was supported by the National Heart, Lung and Blood Institute (NHLBI). RNASeq for "NHLBI TOPMed: Whole Genome Sequencing and Related Phenotypes in the Framingham Heart Study" (phs000974) was performed at the Northwest Genomics Center (HHSN268201600032I), and Genome Sequencing was performed at Broad Genomics (HHSN268201600034I, 3U54HG003067-12S2, 3R01HL092577-06S1). RNASeq for "NHLBI TOPMed: Genetic Epidemiology of COPD (COPDGene)" (phs000951) was performed at the Northwest Genomics Center (HHSN268201600032I), and Genome Sequencing was performed at Broad Genomics (HHSN268201500014C) and the Northwest Genomics Center (3R01HL089856-08S1). RNASeq for "NHLBI TOPMed – NHGRI CCDG: Genes-Environments and Admixture in Latino Asthmatics (GALA II)" (phs000920) was performed at Broad Genomics (HHSN268201600034I), and Genome Sequencing was performed at NYGC Genomics (3R01HL117004-02S3). RNASeq for "NHLBI TOPMed: Study of African Americans, Asthma, Genes and Environment (SAGE)" (phs000921) was performed at Broad Genomics (HHSN268201600034I), and Genome Sequencing was performed at NYGC Genomics (3R01HL117004-02S3) and the Northwest Genomics Center (HHSN268201600032I). RNASeq for "NHLBI TOPMed: SubPopulations and InteRmediate Outcome Measures in COPD Study (SPIROMICS)" (phs001927) was performed at the Northwest Genomics Center (HHSN268201600032I), and Genome Sequencing was performed at Broad Genomics (HHSN268201600034I). RNASeq for "NHLBI TOPMed: MESA and MESA Family AA-CAC" (phs001416) was performed at the Northwest Genomics Center (HHSN268201600032I) and Broad Genomics (HHSN268201600034I), and Genome Sequencing was performed at Broad Genomics (3U54HG003067-13S1, HHSN268201600034I, HHSN268201500014C). RNASeq for "NHLBI TOPMed: Women's Health Initiative (WHI)" (phs001237) was performed at Broad Genomics (HHSN268201600034I), and Genome Sequencing was performed at Broad Genomics (HHSN268201500014C). RNASeq for "NHLBI TOPMed: Lung Tissue Research Consortium (LTRC)" (phs001662) was performed at the Northwest Genomics Center (HHSN268201600032I), and Genome Sequencing was performed at Broad Genomics (HHSN268201600034I). Core support including centralized genomic read mapping and genotype calling, along with variant quality metrics and filtering were provided by the TOPMed Informatics Research Center (3R01HL-117626-02S1; contract HHSN268201800002I). Core support including phenotype harmonization, data management, sample-identity QC, and general program coordination were provided by the TOPMed Data Coordinating Center (R01HL-120393; U01HL-120393; contract HHSN268201800001I). We gratefully acknowledge the studies and participants who provided biological samples and data for TOPMed.
